# Metagenomics of culture isolates and insect tissue illuminate the evolution of *Wolbachia, Rickettsia* and *Bartonella* symbionts in *Ctenocephalides* spp. fleas

**DOI:** 10.1101/2023.02.06.527313

**Authors:** Alexandra Beliavskaia, Kim-Kee Tan, Amit Sinha, Nurul Aini Husin, Fang Shiang Lim, Shih Keng Loong, Lesley Bell-Sakyi, Clotilde K. S. Carlow, Sazaly Abubakar, Alistair C. Darby, Benjamin L. Makepeace, Jing Jing Khoo

**Author notes:** Corresponding author and email address* Jing Jing Khoo. Repositories:* Sequencing reads and genome assemblies produced in study have been deposited in NCBI Genbank under BioProject accession PRJNA909216. Accession numbers for symbiont genomes: *w*CfeF (CP116767), *w*CfeJ (CP116768), *w*Cori (CP116769), *Rickettsia asembonensis* str. Perak (CP116496), pRAS02 (CP116495) and *Bartonella clarridgeiae* str. Perak (CP116497). *Ctenocephalides orientis* mitochondrial assembly was deposited with the accession number OQ383237.

## Abstract

While fleas are often perceived simply as a biting nuisance and cause of allergic dermatitis, they represent important disease vectors worldwide, especially for bacterial zoonoses such as plague (transmitted by rodent fleas) and some of the rickettsioses and bartonelloses. The cosmopolitan cat (*Ctenocephalides felis*) and dog (*C. canis*) fleas, as well as *C. orientis* (restricted to tropical and subtropical Asia), breed in human dwellings and are vectors of cat-scratch fever (caused by *Bartonella* spp.) and *Rickettsia* spp. of the so-called “transitional group”. The latter includes *R. felis* (agent of flea-borne spotted fever) and *R. asembonensis*, an emerging pathogen. The relatively depauperate flea microbiome can also contain arthropod-specific endosymbionts, including a diverse range of *Wolbachia* strains. Here, we present circularized genome assemblies for two *C. orientis*-associated pathogens (*Bartonella clarridgeiae* and *R. asembonensis*) from Malaysia, a novel *Wolbachia* strain (*w*Cori), and the *C. orientis* mitochondrion; all obtained by direct metagenomic sequencing of flea tissues. Moreover, we isolated two *Wolbachia* strains from Malaysian *C. felis* into tick cell culture and recovered circularized genome assemblies for both, one of which (*w*CfeF) is newly sequenced. We demonstrate that the three *Wolbachia* strains are representatives of different major clades (“supergroups”), two of which appear to be flea-specific. These *Wolbachia* genomes exhibit unique combinations of features associated with reproductive parasitism or mutualism, including prophage WO, cytoplasmic incompatibility factors, and the biotin operon of obligate intracellular microbes. The first circularized assembly for *R. asembonensis* includes a plasmid with a markedly different structure and gene content compared to the published plasmid; moreover, this novel plasmid was also detected in cat flea metagenomes from the US. Analysis of loci under positive selection in the transitional group revealed genes involved in host-pathogen interactions that may facilitate host switching. Finally, the first *B. clarridgeiae* genome from Asia exhibited largescale genome stability compared to isolates from other continents, except for SNPs in regions predicted to mediate interactions with the vertebrate host. These findings highlight the paucity of data on the genomic diversity of *Ctenocephalides*-associated bacteria and raise questions regarding how interactions between members of the flea microbiome might influence vector competence.

**Impact statement:** While fleas are vectors for several globally-distributed bacterial zoonoses, including plague, murine typhus, cat-scratch fever, and flea-borne spotted fever, data on the genomic diversity of the etiological agents of these diseases and other members of the flea microbiome are scant. Here, we focused on *Ctenocephalides* spp. fleas associated with dogs and cats in Malaysia, a region where flea microbiome data are particularly sparse. Using a combination of direct metagenomic sequencing of flea tissues coupled with culture isolation and sequencing of infected cell lines, we obtained the first genomes for a cat-scratch agent (*Bartonella clarridgeiae*) and an emerging human pathogen (*Rickettsia asembonensis*) from Asia, as well as genomes for three different *Wolbachia* endosymbiont strains and the first mitochondrial genome for *Ctenocephalides orientis*. The pathogen genomes reveal evidence of selection in genes involved in host-pathogen interactions and in the case of *R. asembonensis*, a novel plasmid genome was obtained. Two of the *Wolbachia* genomes are entirely novel and exhibit highly unusual combinations of features associated with potential reproductive manipulations and mutualistic roles. Our findings highlight major gaps in our understanding of flea microbiome diversity worldwide, with implications for the emergence of bacterial pathogens transmitted by these ubiquitous ectoparasites associated with human dwellings.

**Data summary:** Sequencing reads and genome assemblies produced in study have been deposited in NCBI Genbank under BioProject accession PRJNA909216. Accession numbers for symbiont genomes: *w*CfeF (CP116767), *w*CfeJ (CP116768), *w*Cori (CP116769), *Rickettsia asembonensis* str. Perak (CP116496), pRAS02 (CP116495) and *Bartonella clarridgeiae* str. Perak (CP116497). *Ctenocephalides orientis* mitochondrial assembly was deposited with the accession number OQ383237.

**The authors confirm all supporting data, code and protocols have been provided within 86 the article or through supplementary data files.**

## Introduction

In addition to their direct impact as ectoparasites that can cause flea allergy dermatitis in companion animals and humans, fleas (order Siphonaptera) are major vectors of zoonotic diseases [1]. In contrast with mosquitoes, the public health burden of flea-borne diseases can be attributed almost exclusively to bacterial pathogens rather than viruses or parasites. *Yersinia pestis* (Enterobacterales), the agent of plague, has the greatest current impact worldwide of any flea-borne disease and is one of the most consequential pathogens in human history due to the death toll of the three major plague pandemics. During outbreaks in human populations, plague is transmitted primarily by the Oriental rat flea, *Xenopsylla cheopis*, whereas the human flea, *Pulex irritans*, and fleas of companion animals (*Ctenocephalides* spp.), have been considered as poor plague vectors [2]. A second zoonotic disease transmitted predominantly by *X. cheopis* is murine typhus, caused by *Rickettsia typhi* (Rickettsiales), which is mainly associated with urban or peri-urban environments in the tropics and subtropics [3].

The genus *Ctenocephalides* originated in Africa but now includes two cosmopolitan species, the cat flea *Ctenocephalides felis* and the dog flea *Ctenocephalides canis*, although *C. felis* is more prevalent on dogs worldwide than *C. canis*. The latter is absent from latitudes between 27N and 31S, but in subtropical and tropical Asia, *Ctenocephalides orientis* is prevalent on dogs sympatrically with *C. felis* [4]. *Ctenocephalides* spp. are vectors of *Bartonella henselae* and *Bartonella clarridgeiae* (Hyphomicrobiales) in cats; these pathogens can be transferred to humans by scratches or bites contaminated with flea faeces, causing cat-scratch fever [5]. *Ctenocephalides* spp. are also vectors of several *Rickettsia* spp., including *R. typhi* in some urban centres in the Americas [3]. Another important rickettsial pathogen associated with *C. felis* is *Rickettsia felis*, the agent of flea-borne spotted fever, which is a member of the “transitional group” (TRG) in the rickettsial classification scheme and has a worldwide distribution [6]. Molecular and serological evidence indicates that *R. felis* is a human pathogen capable of inducing severe, potentially fatal, disease [7-10]. While *R. felis* is also a symbiont of non-haematophagous booklice (Psocoptera) and has been detected in many groups of haematophagous arthropods [11], recent data suggests dogs are the main reservoir of infection and *C. felis* is the principal vector [12]. Several organisms closely related to *R. felis* have been detected in *Ctenocephalides* spp. fleas and other arthropods worldwide, including *Rickettsia asembonensis* and *Candidatus* Rickettsia senegalensis, which have been isolated in insect cell culture from *C. felis* specimens [13, 14]. In the case of *R. asembonensis*, there is molecular evidence for human infections from several countries but no indication to date that this species causes severe disease [15]. Earlier surveillance studies have indicated the presence of *Rickettsia felis*-like organisms in *C. orientis* found in the villages of indigenous people in Malaysia [16]; however, these organisms remain uncharacterised.

Few microbiome studies have been performed in fleas, but the evidence to date suggests that in common with other obligate haematophagous arthropods, adult fleas have a simple microbiome [17-20]. In addition to the vectored pathogens listed above, symbionts that are widespread in multiple arthropod taxa are also found in fleas, especially *Wolbachia* (Rickettsiales) and to a lesser extent, *Cardinium* (Sphingobacteriales). These symbiont genera are best known as reproductive parasites that can induce cytoplasmic incompatibility (CI) and other phenotypes that facilitate rapid spread through arthropod populations [21]. However, there is no experimental evidence that flea symbionts are reproductive manipulators [22], although introgression of *C. canis* mitochondrial haplotypes into *C. felis* appears to be linked to *Wolbachia* infection [23]. Fleas are unusual in harbouring diverse *Wolbachia* strains that fall outside the typical clades or “supergroups” infecting ∼50% of all terrestrial arthropods (the so-called pandemic lineages, A and B) [24]. The cat flea genome project, which sequenced a North American *C. felis* lineage, identified two novel *Wolbachia* genomes that were distinct from all known supergroups and included a strain (designated *w*CfeJ) that was hypothesised to induce CI on the basis of its intact *cinAB* operon [25, 26]. The second *Wolbachia* genome (designated *w*CfeT) from North American fleas revealed a capacity to provision biotin, a vitamin that is deficient in blood diets [26].

In the bedbug *Cimex lectularius*, a *Wolbachia* strain from supergroup F has been experimentally demonstrated to provision biotin in an obligate mutualism, which is an unusual phenotype for a *Wolbachia* infection in an arthropod [27]. Recently, we isolated a group F *Wolbachia* (*w*CfeF) from Malaysian *C. felis* into tick cell culture [28] and maintained the strain in a tick cell line; however, since no genome was available, its potential role in vitamin provisioning was unknown. Here we present the genome of this group F *Wolbachia* strain and a second culture isolate related to *w*CfeJ. As part of an effort to isolate and characterise the previously reported *C. orientis*-associated rickettsial organisms from the indigenous communities [16], we also analyse a metagenomic dataset from *C. orientis* for the first time, revealing complete genomes related to *w*CfeT and new strains of *B. clarridgeiae* and *R. asembonensis*. Our data challenge current hypotheses about the roles of *Wolbachia* in *Ctenocephalides* spp. and provide new insights into the evolution of *B. clarridgeiae* and the TRG rickettsiae.

## Methods

### DNA extraction from Wolbachia-infected tick cell cultures

*Wolbachia* isolated from a pool of five *Ctenocephalides* sp. fleas into the *Ixodes scapularis* tick cell line IDE8 [29] was maintained as previously described [28]. Approximately 2.5 years after initial isolation, semi-purified, cell-free *Wolbachia* from infected IDE8 cultures at passage (P)3 was used to infect cells of the *Rhipicephalus microplus* cell line BME/CTVM23 [30]. Semi-purification involved passing the *Wolbachia*-infected IDE8 cells ten times through a 26-gauge needle, pelleting the cell debris by centrifugation at 2,000 g for 5 min and transferring the supernatant onto BME/CTVM23 cells. The infection was maintained by serial passage onto fresh tick cells at intervals. DNA was extracted from the infected parent IDE8 culture, the P4 subculture in IDE8 cells, and a P1 subculture in BME/CTVM23 cells for Nanopore and Illumina sequencing. For DNA extraction, tick cells were detached from the bottom of the culture tube by gentle aspiration using a 2 ml serological pipette. DNA was extracted from 500 μL aliquots of the cell suspension from each culture using a DNA Blood & Tissue Kit (Qiagen) following the manufacturer’s protocol.

### DNA extraction from Ctenocephalides spp. fleas

*Ctenocephalides* spp. fleas were collected from domestic dogs and cats from a village of indigenous people, also known as the Orang Asli, in Perak, Malaysia (4°18′53” N, 100°55′49” E) in August 2019. All field sampling was conducted with the approval of the Universiti Malaya Institutional Animal Care and Use Committee as well as the Department of Orang Asli Development in Malaysia. Live fleas were preserved in 80% ethanol and transported back to the laboratory for storage at 4 °C before further processing. Flea specimens were identified to species level using morphological keys [31]. For DNA extraction, seven to ten individual fleas were pooled according to species and sex, giving a total of 12 pools (named F1-F12, Table S1). DNA was extracted from the pooled flea specimens using a DNA Blood & Tissue Kit (Qiagen) as follows. The exoskeleton of each flea specimen was sliced open with a sterile scalpel. For each pool, the specimens were transferred into a 1.5 mL tube containing the recommended volume of lysis buffer from the kit. For each tube, a sterile pellet pestle was used without the handheld motor to gently grind the specimens and release the tissues into the buffer. The specimens were incubated in the lysis buffer up to three hours and the remaining steps of the DNA extraction were performed according to the kit protocol. Concentration and purity of the extracted DNA samples were determined using a NanoDrop™ 2000 spectrophotometer (Thermo Fisher Scientific).

### Detection of rickettsial DNA from Ctenocephalides spp. fleas

In order to identify *Ctenocephalides* spp. pools with rickettsial infection for metagenomic sequencing, detection of rickettsial DNA from the pooled flea specimens was performed by quantitative PCR (qPCR) amplification of a 76 bp fragment of the rickettsial citrate synthase (*gltA*) gene using previously published primers and protocols [32]. The qPCR was performed using a CFX96 touch real-time PCR detection system (Bio-Rad) with a 6-carboxyfluorescein (FAM) and black-hole quencher (BHQ1-) labelled TaqMan probe (Integrated DNA Technologies). All reactions were prepared in 25 μL reaction volumes, comprising final concentrations of 1 × TaqMan fast advanced master mix (Applied Biosystems), 200 nM of each primer and probe and 1 μL of DNA extracted from the pooled flea specimens.

### Library preparation and sequencing

Prior to library preparation, the concentration of the DNA in each sample was quantified using a Qubit dsDNA BR Assay Kit (Invitrogen). Library preparation for Nanopore sequencing was performed using the protocols for SQK-LSK109 Ligation Sequencing Kit and EXP-NBD104 Native Barcoding Kit (Oxford Nanopore Technologies) to allow for multiplexing. The sequencing libraries were loaded onto MinION Flow Cells (R9.4.1) on a MinION device. Post sequencing, the base-calling of the fast5 reads and demultiplexing were performed using the Guppy base-calling software version 4.2.2 using high accuracy mode, with a quality score cut-off at 9 and minimum read length filter set of 200.

Illumina library preparation and sequencing of the samples were performed in the Centre for Genomic Research (CGR) at the University of Liverpool. For pooled *C. orientis* DNA samples, TruSeq PCR-free paired-end libraries (2×150 bp) with a 350 bp insert were generated and sequenced on a HiSeq 4000. For *Wolbachia* isolates from tick cell culture, NEBNext Ultra II FS kit paired-end libraries (2×150 bp) with a 350 bp insert were generated and sequenced on a NovaSeq 6000 using SP chemistry. The CGR performed the following read curation: the raw fastq files were trimmed for the presence of Illumina adapter sequences using Cutadapt v1.2.1 with option -O 3 [33]; the reads were further trimmed using Sickle v1.200 with a minimum window quality score of 20 (https://github.com/najoshi/sickle); reads shorter than 20 bp after trimming were removed.

### Taxonomic classification and genome assembly

Taxonomic classification of the metagenomic sequencing data was performed using Kraken2 v2.0.7 [34] and the kraken2_nt_20200127 database. Sequence reads classified to *Rickettsiaceae, Wolbachia, Bartonellaceae* or *Anaplasmataceae* were extracted from the Kraken2 output using the extract_kraken_reads.py script from KrakenTools (https://github.com/jenniferlu717/KrakenTools). Bacterial genome assembly from the extracted reads was performed using the long reads assembler Flye v2.8.1 [35]. Genome completeness was assessed by using the Benchmarking Universal Single-Copy Orthologs (BUSCO) pipeline v5.beta.1 and the rickettsiales_odb10 database [36]. The Nanopore-based assemblies were polished with the Nanopore reads using multiple rounds of Racon v1.4.20 [37]. Each round of polishing was evaluated with BUSCO to assess genome completeness. The resulting assembly with the highest BUSCO score was used for multiple rounds of polishing with the corresponding Illumina reads using Pilon v1.24 [38]. Similarly, the polished assemblies were evaluated with BUSCO to assess genome completeness and the assembly with the highest BUSCO was chosen for further analyses.

Publicly available Illumina short reads data (accession: SRX9015637) was used for the assembly of plasmid contigs from the *R. asembonensis* reported from *C. felis* in the north eastern US [39]. The Illumina reads were filtered with BBDuk v26.01.2021 (https://sourceforge.net/projects/bbmap/) to remove adaptor sequences and assembled using metaSPAdes [40].

### Genome annotation and comparison

Gene coding sequences (CDS), ribosomal RNAs (rRNA) and transfer RNAs (tRNA) were annotated from the final assemblies using Prokka v1.13 [41]. Analyses of functional genes was performed using KEGG Automated Annotation Server (KAAS) [42] and KEGG Mapping tools v5.0 [43]. Insertion sequence (IS) elements were determined using the ISsaga web application pipeline [44]. IS elements flagged as probable false positives were excluded from the dataset. Genome wide variant calling was performed using SNIPPY v4.6.0 (https://github.com/tseemann/snippy) and mapping the corresponding Illumina reads to the assembled genomes. Comparisons of closely related genomes were performed and visualised using dot-plots produced by the D-GENIES web tool based on a Minimap2 alignment [45, 46] and Circos representations of progressiveMauve v2.4.0 [47] alignments produced using Circos v0.69.8 [48, 49] implemented in the Nanogalaxy platform [50]. All genomes were rotated to start at *dnaA* prior to synteny analyses. Additional gene prediction and annotations for the *Bartonella clarridgeiae* assembly were obtained from the comprehensive genome analysis pipeline implemented in PathoSystems Resource Integration Center (PATRIC) web portal v3.6.12 [51], which provides RAST tool kit (RASTtk)-based annotation [52] and more functional information for pathogens.

### Phylogenomic tree construction

Ortholog sequences were produced using Orthofinder v2.5.4 [53]. Multiple sequence alignments of the protein sequences for each single copy orthogroup (OG) were performed using MAFFT v7.149bb [54]. Noisy or poorly aligned protein positions within each protein block were trimmed using Gblocks v0.91b [55]. The trimmed blocks were concatenated into a supermatrix for constructing the maximum likelihood trees using IQTREE v2.1.2 [56]. Separate models appropriate for each protein in the supermatrix were determined by ModelFinder [57] within IQTREE. Branch support was calculated using the following options in IQTREE: (i) ultra-fast bootstrap, (ii) SH-aLRT support, (iii) local bootstrap support and (iv) aBayes bayesian support, with all options set to 1,000. All options produced highly similar values for each branch and the values from the ultra-fast bootstrap option [58] were displayed along with the consensus trees in the final figures. The Interactive Tree of Life online tool was used to visualise the consensus trees produced (https://itol.embl.de).

### Analyses of Wolbachia prophage regions, cytoplasmic incompatibility factor (cif)-like genes, biotin operon and plasmid region

Prophage regions in *Wolbachia* assemblies were estimated using the PHASTER web application tool [59]. CDS flanking or located within the PHASTER-predicted prophage regions were subjected to BLASTp queries using a previously published dataset for *Wolbachia* phage (WO) annotations [60] and the NCBI non-redundant protein database to identify the genes in eukaryotic association modules (EAM). Bayesian-inferred phylogeny for *Wolbachia* phage large serine recombinase was produced using MrBayes v3.2.7a [61] with two runs, four chains and 1,000,000 generations. ProtTest v3.4.2 was used to select for best protein model [62].

To determine the presence of *cif*-like gene homologues, amino acid sequences translated from the nucleotide sequences of *cif*-like genes identified in a previous study [63] were used in BLASTp queries of a database of the CDS from the *Wolbachia* assemblies from this study or other published *Wolbachia* genomes. Protein domains within the *cif*-like homologs and EAMs were determined using the HMMR search in the Simple Modular Architecture Research Tool (SMART) web platform v9 [64] and manual inspection of the aligned amino acid sequences of closely related *cif*-like genes obtained using Clustal Omega [65]. Visualisation of CDS in *Wolbachia* prophage regions and protein domains within the *cif*-like genes was performed using the Illustrator of Biological Sequences software v1.0.3 [66]. Comparison of *Wolbachia* biotin operons and plasmid regions was performed using BLASTn and visualised using GeneplotR v0.8.11 [67].

### Positive selection analysis for TRG Rickettsia

To produce codon-aware alignments, nucleotide sequences corresponding to the single copy OGs from TRG *Rickettsia* were aligned individually using PRANK v170427 [68] and concatenated. Maximum likelihood phylogeny was produced from the concatenated nucleotide alignment with IQTREE v2.0.3 [56] with 1,000 ultra-fast bootstraps and best codon model selection from ModelFinder [57]. Vertebrate pathogens were marked as foreground branches using the web-based phylotree widget (http://veg.github.io/phylotree.js/). Signals of positive selection were determined using the aBSREL branch-site model in HYPHY v2.5.39 [69] with the codon-aware alignment for the individual OGs and the maximum likelihood tree produced. OGs with signals of positive selection in at least one branch (P < 0.05 after correction for multiple testing) were submitted to the eggnog-mapper v2 web service for functional annotation [70].

### Assembly, annotation and phylogenomic analyses of C. orientis mitochondrial genome

Nanopore reads from *C. orientis* F5 pool were aligned against *C. felis* EL2017-DRISC mitochondrial assembly (accession: MT594468) using minimap2 v2.24-r1122 [46] and extracted into a separate file using samtools v1.15. [71]. Filtlong v0.2.1 (https://github.com/rrwick/Filtlong) was used to choose the best reads constituting 2 Mb corresponding to approximately 100X coverage of the mitochondrial genome of *C. felis*. These reads were assembled with Flye v2.8.3-b1695 [35]. The assemblies were polished with Illumina reads using nthits v0.1.0 with options -b 36 -k 40 and ntedit v1.3.5 [72] in ‘best substitution or best indel’ mode (-m 1). The assemblies were rotated to start with the *nad2* gene. The MITOS2 webservice [73] with the RefSeq 89 Metazoa reference and genetic code 5 for Invertebrates was used for the annotation.

Eight mitochondrial assemblies for Siphonaptera were used for the phylogenetic reconstruction, while three non-siphonapteran assemblies (*Broscus cephalotes, Euclimacia badia* and *Abachrysa eureka*) were used as an outgroup. Mafft v7.480 [54] was used for sequence alignment of 13 protein-coding and two ribosomal genes (Table S2), whereas BMGE v1.12 [74] was used to select phylogenetically informative regions from the multiple sequence alignments. Modeltest-ng [75] was used to estimate the best substitution model for each gene, and IQ-TREE v.2.1.4-beta [56] was used for phylogenetic tree reconstruction with 1,000 bootstraps. Each of the protein-coding and rRNA genes was treated as a separate partition (Table S2). The resulting phylogenetic tree was visualized using Interactive Tree of Life v4 (https://itol.embl.de). BLAST Ring Image Generator (BRIG) [76] was used for comparison of closely related mitochondrial genome assemblies. Clinker v0.0.28 [77] was used for visualisation of gene order in siphonapteran mitochondrial assemblies.

## Results

### Metagenomic assemblies from wCfeF-infected tick cell cultures and from C. orientis

During the first attempt at Nanopore sequencing of the parent *Wolbachia*-infected IDE8 culture, Flye assembly using reads assigned to *Anaplasmataceae* from Kraken2 revealed the presence of two circularised contigs at 55x and 25x coverage respectively (Fig. S1a). Both contigs had different sizes (∼1.4 mb vs ∼1.2 mb) and genome alignment also suggested very limited similarity and synteny between these two genomes (Fig. S1d). This finding provided more evidence for the previously-suspected presence of a second *Wolbachia* strain at a lower density than *w*CfeF in the parent IDE8 culture [28]. In an attempt to improve the growth of the putative second strain, the *C. felis*-derived *Wolbachia* was passaged onto fresh IDE8 cells and into a second tick cell line, BME/CTVM23, previously shown to be susceptible to a range of *Wolbachia* strains [28, 78]. DNA extracts from P4 IDE8 and P1 BME/CTVM23 subcultures were used for the subsequent round of Nanopore and Illumina sequencing. Both genomes were found in the Flye assemblies from the infected IDE8 (Fig. S1b) and BME/CTVM23 (Fig. S1c) subcultures. The Nanopore fastq datasets from all three *Wolbachia*-infected IDE8 (parent and subculture) and BME/CTVM23 cell cultures were combined to improve coverage, which yielded two complete circularised genomes at 141x and 87x coverage, respectively (Table 1). The resulting Flye assemblies were polished with the combined Illumina dataset from *Wolbachia*-infected subcultures of IDE8 and BME/CTVM23. Comparison of the *wsp* gene sequence from the larger assembly (∼1.4 mb) confirmed that it represents *w*CfeF, the strain previously described from the infected parent IDE8 cell culture [28].

**Table 1.**
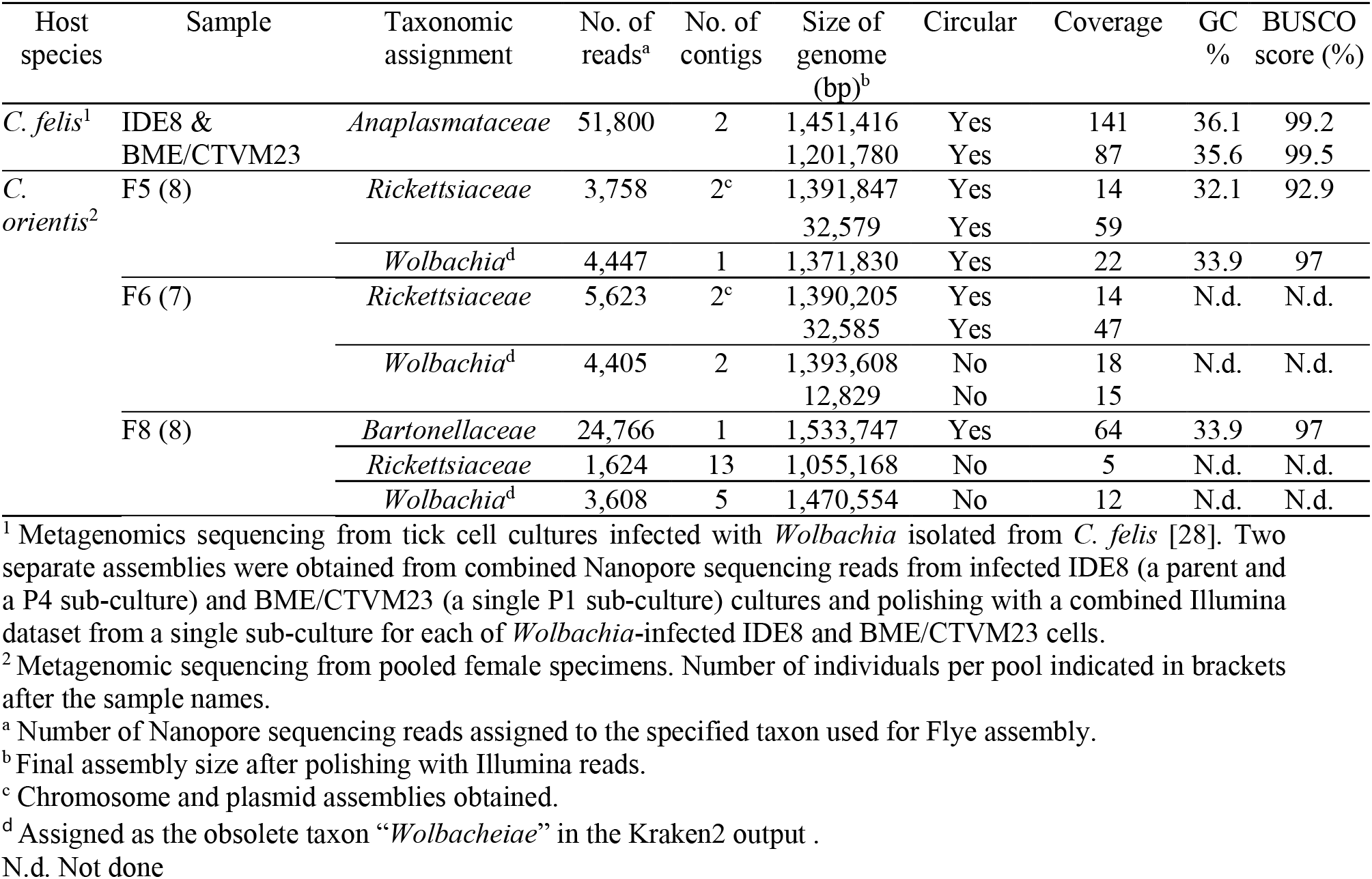
Metagenomic assemblies obtained from *Ctenocephalides orientis* and from *Wolbachia*-infected tick cell cultures

Quantitative PCR amplification of the rickettsial *gltA* gene detected the presence of rickettsial DNA in all *C. orientis* pools (Ct values ranging from 23.12 to 26.28, Table S1) but not in the *C. felis* pools. Three pools (F5, F6, and F8) of female *C. orientis* were selected for metagenomic sequencing in order to recover rickettsial genome sequences; however, reads assigned to *Wolbachia* and *Bartonellaceae* taxa were also recovered in addition to the *Rickettsiaceae* taxon (Table 1). Final assemblies were obtained from the F8 pool from these reads (Table 1) for all three taxa; however, only the *Bartonellaceae* assembly was circularised. While complete circularised *Rickettsiaceae* chromosomal and plasmid assemblies were obtained from both the F5 and F6 pools, only draft assemblies with 13 contigs were obtained for *Rickettsiaceae* from the F8 pool. Genome alignments comparing the *Rickettsiaceae* assemblies from F5 and those from F6 and F8 showed high identities (Fig. S2); therefore, only the *Rickettsiaceae* assemblies from F5 were used for further analyses below. A complete circularised assembly for *Wolbachia* was only obtained from the F5 pool. Draft assemblies from the F6 and F8 pools were obtained in two and five contigs, respectively. Genome alignment comparing the *Wolbachia* assemblies from F5 and those from F6 and F8 also showed high identities between the assemblies (Fig. S2). The assembly from F5 was named *Wolbachia* endosymbiont of *C. orientis* (*w*Cori) and used for further analyses below. The high BUSCO scores (92.9% - 99.5%, Table 1) indicate the high level of completeness of the genomes assembled in this study.

### Wolbachia genomes from Ctenocephalides spp. represent novel supergroup V and additional members in supergroups I and F

Consistent with earlier multi-locus sequence typing data [28], *w*CfeF was found in the same clade as the other supergroup F members in phylogenetic analyses (bootstrap support 100, Fig. 1a). Moreover, *w*CfeF formed a sister clade to *w*Cle and *w*Mhie (bootstrap support 100, Fig. 1b) in an additional phylogenetic analysis performed for supergroup F members only. The smaller *Wolbachia* assembly (∼1.2 mb) clustered with the *w*CfeJ isolate from the *C. felis* colonies in the US [25, 26], forming a clade independent of supergroups F and S with robust support (bootstrap support 100), thereby providing evidence for a novel supergroup [designated as V (Fig. 1a)]. Similarly, *w*Cori and *w*CfeT clustered together independently from *w*Fol (supergroup E) and *w*Pni (supergroup M) (bootstrap support 93, Fig. 1a). We assign these genomes to the existing supergroup I on the basis of similarity to published *Wolbachia* 16S sequences from fleas (Fig. S3). Finally, our analysis placed *w*How (hosted by the tylenchid nematode, Howardula sp.) from a recent study as the sole known member of a new supergroup, consistent with published findings [79]. We designate the *w*How clade as group W.

**Fig. 1.**
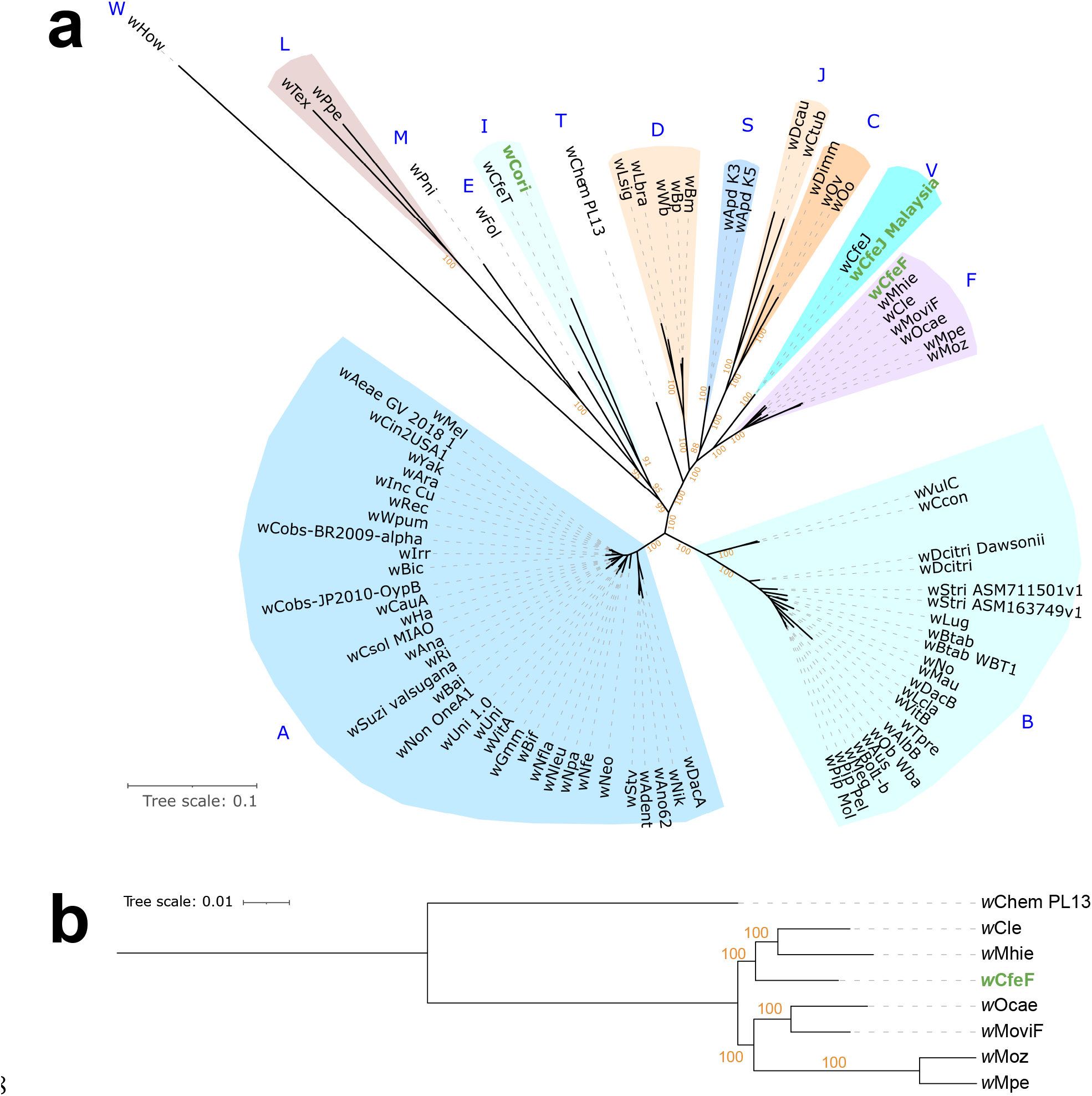
(a) Maximum-likelihood phylogeny based on the concatenated alignments of 45 single copy orthologs across 86 *Wolbachia* genomes (7,110 amino acid sites,) using a partitioned best-fit model for each ortholog. (b) Maximum-likelihood phylogeny of supergroup F *Wolbachia*, with *w*Chem PL13 as outgroup. The tree is based on the concatenated alignments of the 500 single copy orthologs (147,887 amino acid sites) with the best-fit models partitioned fit to each ortholog. The trees are rooted at mid-point. The newly assembled genomes are marked in green.

### Wolbachia genomes recovered from Ctenocephalides spp

The *w*CfeF genome comprised 1,451,416 bp in a single contig with a GC value of 36.1%. Prokka prediction produced 1,359 CDS, 34 tRNAs and three rRNAs. Comparison to the *w*Cle genome (from *C. lectularius*) revealed limited similarity and synteny between these two genomes (Fig. 2a). ISSaga predicted 65 IS elements from 12 different families across the genome at 47.61-100% similarities. The positions of many of the IS elements appeared to correspond with regions on the *w*Cle alignment where breaks in synteny occurred (Fig. 2a), suggesting that IS elements may have contributed to the genomic rearrangements between these F-group *Wolbachia* genomes. Three prophage regions were found within the *w*CfeF genome, which were absent from the *w*Cle genome (Fig. 2a).

**Fig. 2.**
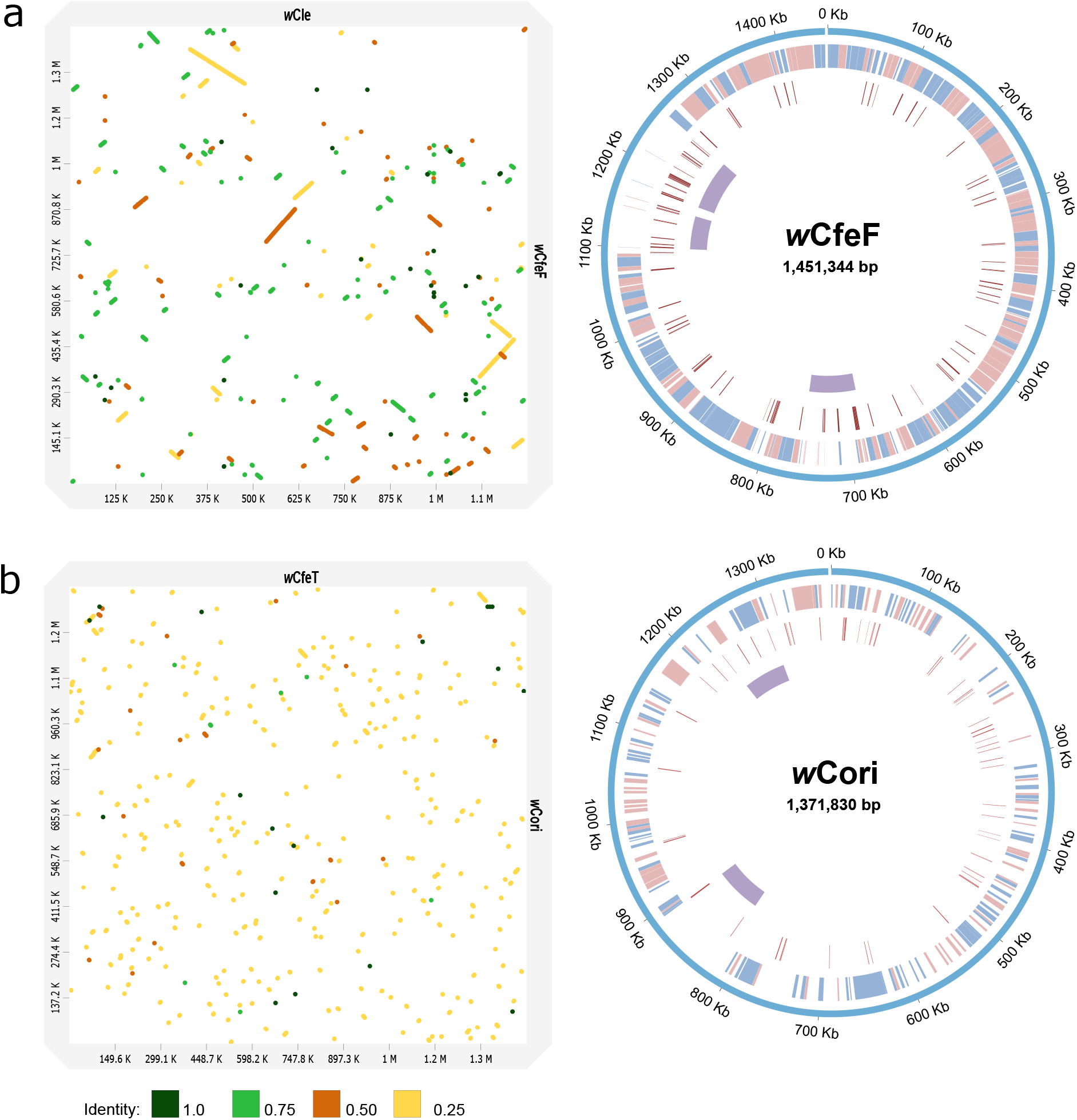
*Wolbachia* genome comparisons. Left panels: Minimap2 alignment of (a) *w*CfeF and *w*Cle (accession: NZ_AP013028.1), and (b) *w*Cori and *w*CfeT (accession: NZ_CP051156.1). The identity values refer to a BLAST-like alignment identity computed from the alignment file and have been binned into four groups (<0.25, 0.25 – 0.5, 0.5 – 0.75, 0.75 – 1). Right panels: Circos representation of (a) *w*CfeF and (b) *w*Cori genomes. Outer ring: (a) *w*CfeF and (b) *w*Cori genomes. First inner ring: locally colinear blocks identified in progressiveMauve alignment between (a) *w*CfeF and *w*Cle, and (b) *w*Cori and *w*CfeT genomes. Second inner ring: IS or other mobile genetic elements predicted in each genome. Third inner ring: predicted prophage regions in each genome. The colour blue represents direct matches and red represents reverse matches in the first inner ring.

The current *w*CfeJ genome was 1,201,780 bp in size, with a GC value of 35.6%. Genome comparison showed high level of similarity and synteny with the *w*CfeJ assembly from the US (accession: NZ_CP051157.1, Fig. S4). Compared to this North American genome [25, 26], only 18 nucleotide variants were detected in the Malaysian *w*CfeJ genome (Table S3), indicating that both genomes are highly similar to each other. Nine variants were due to non-synonymous substitution, which were found in the gene for elongation factor Tu (*tuf*), DEAD/DEAH box helicase, CvpA family protein, ABC transporter protein and four hypothetical proteins; whereas two frameshift mutations were observed in hypothetical proteins. The remaining variants were due to synonymous substitutions. PHASTER prediction did not reveal the presence of phage elements in our *w*CfeJ genome; however, 52 IS were predicted (47.61 – 100% similarity).

The genome of *w*Cori is 1,371,830 bp in size with a GC value of 33.9%. Alignment to the *w*CfeT genome indicated limited synteny between these two genomes (Fig. 2b). A total of 62 IS elements from 14 different families (48.74 – 100%) were found in the genome, with the majority from the IS110 family. Similar to *w*CfeF, the majority of the IS appeared to be found in regions on the *w*CfeT alignment where breaks in synteny occurred. We identified two prophage regions in the genome, with one prophage region sharing similarities with a prophage sequence in *w*CfeT. Prokka prediction produced 1,610 CDS in *w*Cori, with one tmRNA, 34 tRNAs and three rRNAs.

### Multiple prophage variants found in Wolbachia from Ctenocephalides spp

Prophage regions in *w*CfeF and *w*Cori were identified by PHASTER and by performing BLASTp queries using the Prokka-annotated CDS flanking the PHASTER-predicted regions. In addition, the recently available data for *Wolbachia* phage (WO) annotations [60] as well as the non-redundant protein sequence database in NCBI GenBank were used to facilitate prophage annotations. We found three prophage regions in *w*CfeF that were complete, with the core phage modules having sizes of 78,289 bp (WOCfeF1), 78,066 bp (WOCfeF2) and 84,737 bp (WOCfeF3), respectively (Fig. 3a). WOCfeF2 and WOCfeF3 formed a continuous region with two sets of core phage modules. Two prophage regions with all core phage modules were also found in *w*Cori, and were 74,721 bp (WOCori1) and 66,861 bp (WOCori2) in length, respectively. However, a number of genes within the core phage modules were found to be pseudogenised and thus it is unclear if these prophage regions could enter a lytic cycle. BLASTp analyses also revealed many CDS across *w*CfeF and *w*Cori genomes matched to phage-associated genes in the WO database, even though the CDS were not located within the prophage regions with complete core phage modules.

**Fig. 3.**
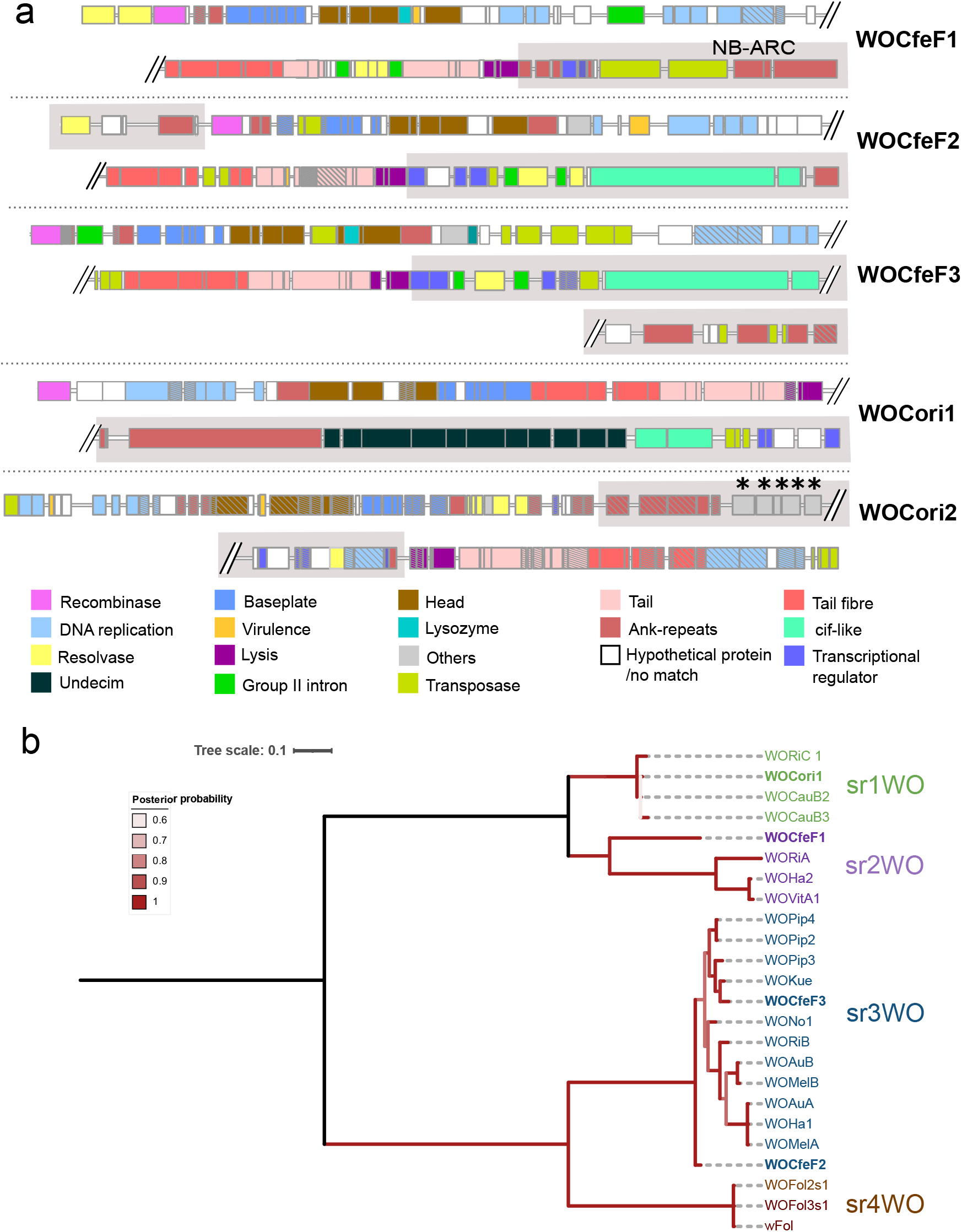
(a) Prophage regions with core phage modules in *w*CfeF and *w*Cori genomes. Eukaryotic association module (EAM)-like regions are highlighted in the grey boxes. Pseudogenised genes are marked with diagonal lines. Asterisks indicate the genes shared only between *w*Cori and *w*CfeT, but not other published *Wolbachia* strains. (b) Bayesian phylogeny based on the large serine recombinase in *Wolbachia* prophage regions. Phylogeny was generated from 299 amino acid positions using the JTT + IG model. The tree was rooted at mid-point.

In terms of core module synteny, the prophage regions from *w*CfeF were similar to each other (*i*.*e*., in the order of baseplate, head, DNA replication and repair, tail). The prophage regions in *w*CfeF also had multiple copies of resolvases/recombinases, and some appeared to be associated with group II introns. However, the core modules in WOCori1 and WOCori2 had different arrangements. The large serine recombinase has been recently proposed for WO typing and classification, and four distinct WO variants (sr1WO-sr4WO) have been described based on the large serine recombinase phylogeny and core module synteny [60]. WOCfeF2 and WOCfeF3 clustered with members of the sr3WO members, which share similar core module synteny [60], in a single clade (posterior probability 1) (Fig 3b). WOCfeF1 did not cluster with WOCfeF2 or WOCfeF3; instead, it formed a sister clade with the members of the sr2WO group (posterior probability 1). WOCori1 clustered with sr1WO members (posterior probability 1), which shared similar core module synteny (DNA replication and repair, head, baseplate, tail) and the presence of single domain HTH_XRE transcriptional regulators at the 3’ end of the EAM-like region [60]. WOCori2 was not included in the phylogenetic analysis since the recombinase gene was fragmented.

Regions similar to the EAMs previously reported in other *Wolbachia* strains were also observed in *w*Cori and *w*CfeF prophage regions. The EAM features genes encoding transcriptional regulators and proteins with eukaryotic-like domains, and could potentially contribute to the manipulation of host processes [80]. Many of the CDS within the EAM-like regions found in the prophage sequences here have BLASTp matches to homologs from other published *Wolbachia* phages (Table S4). A protein with ankyrin repeats and the NB-ARC domain was encoded by a CDS in the WOCfeF1 EAM-like region. Cytoplasmic incompatibility factor (*cif*)-like genes, which are associated with host reproductive phenotypes in some insect hosts and previously reported in supergroups A, B, F and T *Wolbachia* genomes [60, 63], were found in the EAM-like regions of WOCfeF2, WOCfeF3 and WOCori1 (see below). WOCori1 also contained a recently-described Undecim Cluster found in supergroup A, B, E and M *Wolbachia* genomes, consisting of a series of 11 genes encoding proteins with a variety of metabolic or transport functions [60]. The EAM-like region of WOCori2 contained five genes that were found in *w*CfeT, comprising genes involved in pantothenate biosynthesis (see section on metabolic pathways).

### cif-like genes identified in Wolbachia from Ctenocephalides spp

Recent phylogenetic studies classified *cif*-like genes into five different types [63]. Two sets of *cif*-like genes (*w*CfeF_1 and *w*CfeF_2) were identified in *w*CfeF and both were associated with the prophage regions WOCfeF2 and WOCfeF3, respectively. The *cif*-like genes in *w*CfeF were found to be more similar to the type V *cif*-like genes (Fig. 4), with the presence of an OTU-like cysteine protease and many ankyrin repeats in the *cif*B homologs. However, the latrotoxin C-terminal commonly associated with type V *cif*B homologs was absent from *w*CfeF_1 and *w*CfeF_2 *cif*B homologues. A single intact set of *cif*-like genes (*w*Cori_2) with 99-100% identity to the type III *cif*-like genes in a WO-like island in *w*VitA was found in *w*Cori within the EAM of the WOCori1 prophage region. Another two sets of highly pseudogenised *cif*-like genes were also found outside the predicted prophage regions. One set (*w*Cori_1) was related to the type I homologues, which are characterised by the presence of a deubiquitinase and peptidase domain in the *cif*B homolog. The final set (*w*Cori_3) was related to type V; however, the *cif*B homolog was highly fragmented. Only the PDDEXK domains and no other domains common to type V homologs were detected in the fragments. The *cif*B homologues in *w*Cori and *w*CfeF are possibly undergoing degradation from larger, modular *cif*B-like genes similar to the *cif*-like genes in *w*CfeJ [26].

**Fig. 4.**
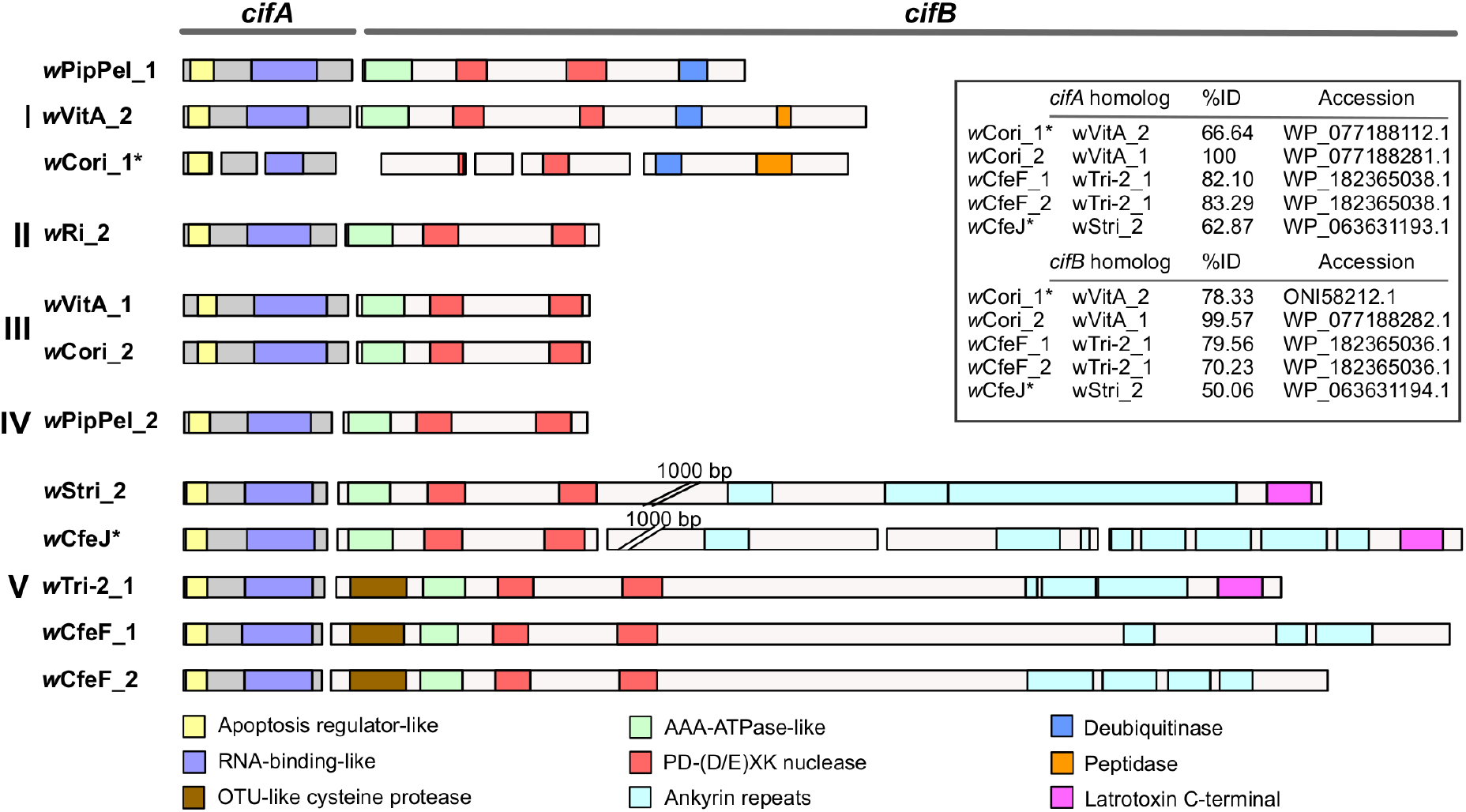
*cif*A/B homologs in *Ctenocephalides* spp.-associated *Wolbachia*. Inset: Nearest BLASTp matches for the *cif-*like genes identified in *Wolbachia* associated with *Ctenocephalides* spp. Asterisks indicate amino acid translation from pseudogenised genes. *w*Cori_3 is not shown here due to its highly fragmented nature.

### Metabolic pathways in Wolbachia from Ctenocephalides spp

Functional analyses by KEGG showed most metabolic pathways, apart from vitamin biosynthesis, were conserved within the *Wolbachia* strains from *Ctenocephalides* spp. (Fig. S5). So far, *w*Cle is the only member of supergroup F known to harbour a complete biotin operon (known as the Biotin synthesis Operon of Obligate intracellular Microbes or BOOM) [81], supporting its role as a nutrition-provisioning symbiont in the bedbug host [27]. *w*CfeF also carried BOOM; however, the biotin synthase (*bioB*) gene was pseudogenised, as confirmed by PCR and Sanger sequencing. The BOOM was absent in *w*Cori, despite its close relationship to *w*CfeT, the other *Wolbachia* from *C. felis* reported to harbour BOOM [26]. *w*Cori and *w*CfeT both contained the genes *panB, panC, panD* and *panG*, which are related to pantothenate biosynthesis. These genes, except *panD*, have only previously been reported in the *Wolbachia* endosymbiont of *Menacanthus eurysternus* (*w*Meur1), a supergroup F strain with a highly reduced genome [82]. As mentioned above, the pantothenate biosynthesis island was found in the EAM region of WOCori2 prophage (Fig. 3a), suggesting a possible role of phages in the lateral gene transfer (LGT) of these genes.

### Biotin operon and plasmid region in Wolbachia from Ctenocephalides spp

Phylogenetic analysis indicated that the *w*CfeF BOOM is more closely related to those of *w*Cle and the other *Wolbachia* associated with *Cimex* sp. (str. KTCN) than to that of *w*CfeT (Fig. 5a), suggesting that BOOM from *w*CfeF and *w*Cle likely share similar origins despite being hosted in different insect orders. Consistent with an earlier study [26], the genes flanking the biotin operons were also dissimilar between *w*CfeF, *w*Cle, and *w*CfeT, indicating separate LGT events as the origin of the biotin operons. Interestingly, a plasmid-like region was found close to the biotin operon in *w*CfeF (Fig. 5b). A recent study described the presence of two circular extrachromosomal plasmid elements, pWALBA1 and pWALBA2, from *Wolbachia* strain *w*AlbA [83]. BLASTp analyses of the proteins in the *w*CfeF plasmid region showed high similarity to the proteins in pWALB1 (percentage identity 49% - 85%, Table S5). The only pWALB1 proteins not found in the *w*CfeF plasmid region were the *parA* and *parG*-like plasmid partition proteins. Comparison of the gene arrangements also revealed high synteny between the *w*CfeF plasmid region and pWALB1 (Fig. 5b). Gene islands containing plasmid-like elements were found in other *Wolbachia* strains, including *w*Cle [83], and inspection of the genes flanking the *w*Cle biotin operon revealed the presence of a plasmid gene island containing hypothetical proteins and a transposon (Fig. 5b).

**Fig. 5.**
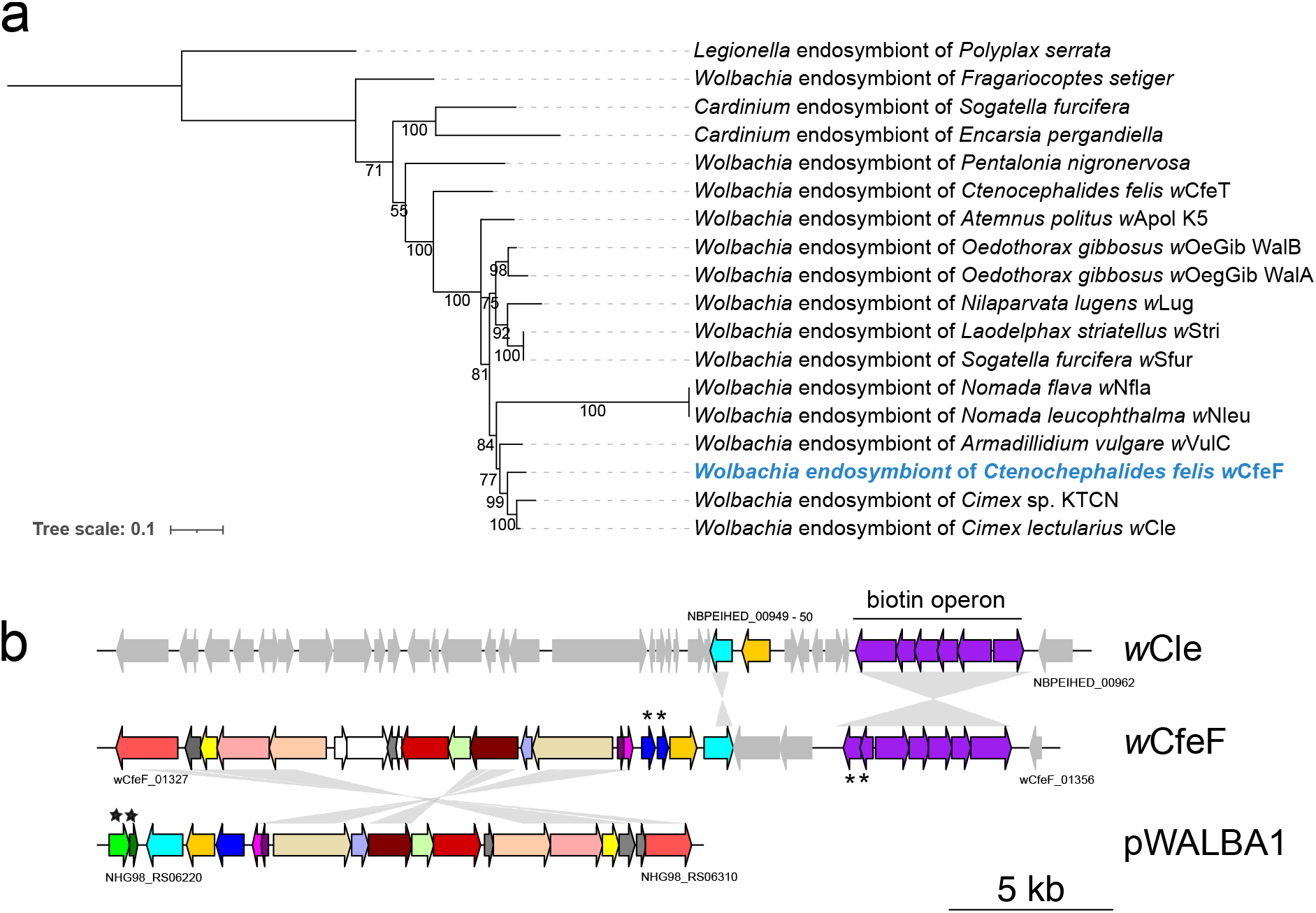
(a) Maximum-likelihood phylogeny based on the concatenated alignments of the proteins from the biotin operon (2,023 amino acid sites, at least four proteins in the operon for each bacterium strain included) of selected strains of *Wolbachia* and *Cardinium*, with a *Legionella* endosymbiont as an outgroup, using JTTDCMut+F+I+G4 as the best-fit model according to BIC. (b) Comparison of biotin operon and plasmid gene organisation in *w*Cle, *w*CfeF, and pWALBA1. Pseudogenised genes are labelled with asterisks. *ParA* and *parG*-like plasmid partition proteins are labelled with stars. BLASTp matches of proteins in *w*CfeF plasmid region to pWALBA1 are given in Table S5. Grey areas between sequences indicate BLASTn matches (E-value < 1e-5).

### Complete genome of R. asembonensis str. Perak

The phylogeny of TRG rickettsiae showed clustering of the current *Rickettsia* genome with *R. asembonensis* NMRCii (bootstrap value 100, Fig. 6a). A maximum-likelihood tree based on three separate genes, *gltA, sca4* and *ompB*, also confirmed the placement of the current *Rickettsia* genome in a clade with various *R. asembonensis* strains (bootstrap value 100, Fig. S6). This genome represents the first for *R. asembonensis* from Asia (named str. Perak). and notably, the only other rickettsial assembly reported from Malaysia (*Rickettsia* sp. TH2014) [14] was more closely related to *Candidatus* R. senegalensis than to *R. asembonensis* (bootstrap value 100, Fig. S6).

**Fig. 6.**
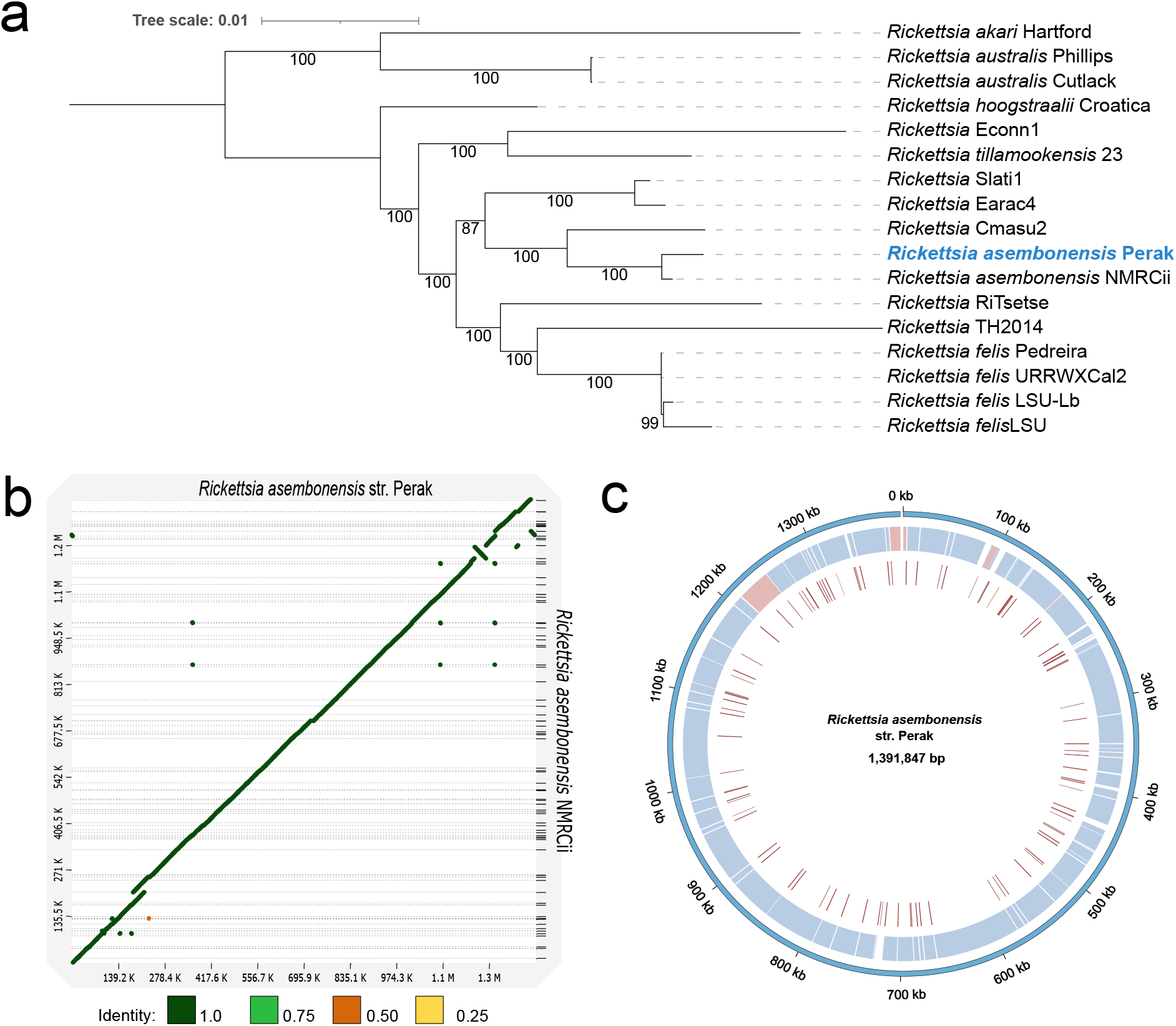
(a) Maximum-likelihood tree of TRG rickettsiae based on 471 single copy orthogroups (OGs) (111,864 amino acid positions) using a partitioned best-fit model for each OG. Bootstrap values above 80 were displayed. Tree was rooted at mid-point. Accession numbers for rickettsial strains within the transitional group used in phylogenetic analysis are given in Table S7. (b) Minimap2 alignment of *Rickettsia asembonensis* str. Perak chromosome and *Rickettsia asembonensis* NMRCii assemblies (accession: GCF_000828125.2). The identity values refer to a BLAST-like alignment identity computed from the alignment file and have been binned into four groups (<0.25, 0.25 – 0.5, 0.5 – 0.75, 0.75 – 1). (c) Circos representation of genomic comparison between *Rickettsia asembonensis* NMRCii and *R. asembonensis* str. Perak chromosome. First inner ring: locally colinear blocks identified in progressiveMauve alignment between *R. asembonensis* str. Perak and the contigs from the *R. asembonensis* NMRCii assembly. Second inner ring: IS or other mobile genetic elements predicted in *R. asembonensis* str. Perak. The colour blue represents direct matches and red represents reverse matches in the first inner ring.

The complete genomic assembly of *R. asembonensis* str. Perak comprised two contigs representing the chromosomal and plasmid sequences. The larger chromosomal contig was 1,391,847 bp in size with a GC value of 32.14% and BUSCO score of 95.9%. Prokka prediction produced 1,553 coding sequences (CDS) from the chromosome, with 917 annotated as hypothetical proteins, 34 tRNAs and three rRNAs. Mapping of the chromosomal contigs from the *R. asembonensis* NMRCii (accession: GCF_000828125.2) assembly to the current genome showed a high level of similarity and synteny between them (Fig. 6b,c). Most of the IS elements predicted in *R. asembonensis* str. Perak appeared to correspond with the missing regions in the *R. asembonensis* NMRCii assembly. One hundred IS elements were predicted with 48.75-100% similarities to previously reported IS elements. As expected for a rickettsial species, *R. asembonensis* str. Perak did not have many genes involved in biosynthetic pathways (Fig. S5). Genome-wide variant analysis using *R. asembonensis* NMRCii as a reference revealed a small number of variants arising from 33 SNPs, four insertions and one deletion in the chromosomal contig (Table S6), with 22 of these resulting in non-synonymous substitution or causing a frameshift. Non-synonymous substitutions were found in coding sequences from bacterial transport systems (TolC, MFS, ABC) and a RelE/ParE family toxin.

### Lack of complete Rickettsiales-amplified gene element in R. asembonensis plasmids

The plasmid assembly from *R. asembonensis* str. Perak was named pRAS02, following the first plasmid described for *R. asembonensis* NMRCii, pRAS01. The top three matches from a BLASTn search with pRAS02 included *Rickettsia raoultii* strain Khabarovsk plasmid pRra3 (query cover [QC]: 23%, percentage identity, PI: 92.5%), *R. asembonensis* NMRCii plasmid pRAS01 (QC: 23%. PI: 85.5%) and *Rickettsia bellii* AN04 unnamed plasmid (QC: 19%, PI: 83.2%). This low QC indicated substantial dissimilarity overall between pRAS02 and pRAS01. Thirty-two CDS were identified from pRAS02, including 12 amino acid sequences with BLASTp matches to other known plasmid proteins and seven with matches to hypothetical proteins (Fig. S7). Most of the remaining CDS were associated with mobile genetic elements, although one did not match any existing entry from the NCBI non-redundant protein database.

Alignment of pRAS02 and the plasmids from the top three BLASTn matches showed limited shared regions (Fig. 7). The *Rickettsiales*-amplified gene element (RAGE) consists of mobile genetic elements previously reported in the chromosomes and/or plasmids of a number of rickettsial species, including *Rickettsia buchneri, R. bellii, Rickettsia peacockii, Rickettsia massiliae, R. felis* and *Rickettsia tamurae* [84-86]. An intact RAGE was not observed in any of the plasmids analysed here; however, a small number of RAGE-associated genes were detected. Furthermore, F plasmid-like Type IV secretion system (T4SS) proteins were mostly absent from the analysed plasmids, except for the presence of TraC, TraV and TraB in the unnamed plasmid from *R. bellii* AN04 (TraC was pseudogenised), whereas TraG was present in pRa3. The *Agrobacterium tumefaciens* Ti plasmid-like T4SS proteins, TraAI (mobilisation A/L and relaxase domains-containing protein) and TraD [84], were also detected in pRa3 (both genes) and the *R. bellii* unnamed plasmid (TraAI only), although they appeared to be pseudogenised in pRAS01 and pRAS02. Moreover, manual inspection of the *R. asembonensis* str. Perak chromosome revealed the presence of proteins normally flanking RAGE, including pseudogenised forms of TraAI, SpoT synthetase/hydrolase, histidine kinase, TraD and citidylate kinase. The tRNA^Val-GAC^ gene, which serves as an insertion site for RAGE [84], was also found. BLASTn analyses did not find any plasmid CDS in the chromosome.

**Fig. 7.**
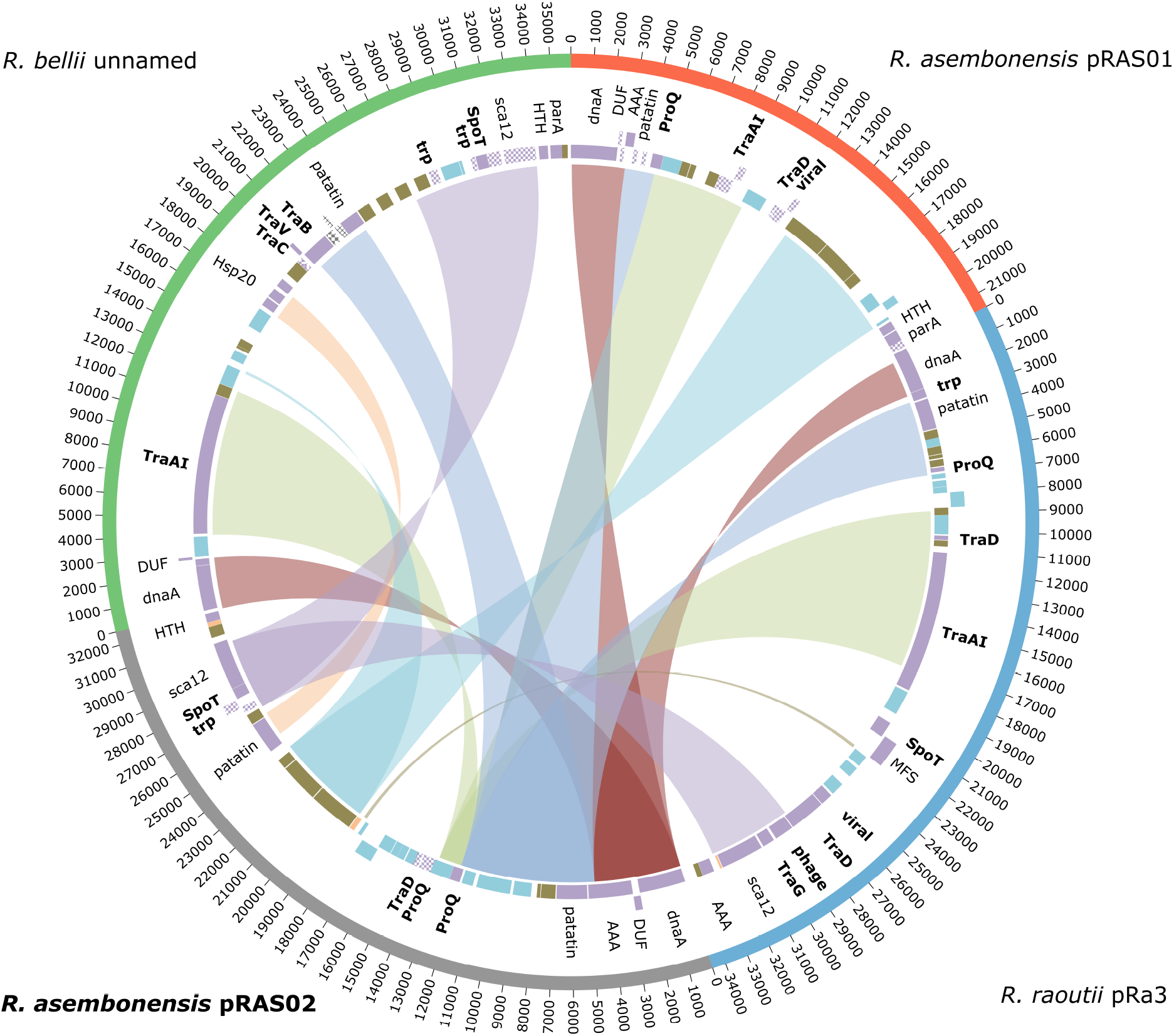
Circos representation of the progressiveMauve alignment of *Rickettsia asembonensis* str. Perak plasmid pRAS02, *Rickettsia raoultii* strain Khabarovsk plasmid pRra3 (accession: NZ_CP010972.1), *R. asembonensis* NMRCii plasmid pRAS01 (accession: NZ_CP011517.1) and *Rickettsia bellii* AN04 unnamed plasmid (accession: NZ_CP015011.1). Genes in bold represent genes associated with RAGE.

We attempted to assemble plasmid sequences from a previously reported North American *R. asembonensis* genome using the publicly available metagenomic short reads data (accession: SRX9015637) obtained from infected *C. felis* [39]. The assembly produced two contigs, of 36,703 bp and 7,177 bp respectively, which were aligned to pRAS02 (Fig. S8a), suggesting the presence of plasmids in the US *R. asembonensis* genome. However, it is unclear if both the contigs in the US genome represent the complete plasmid since they were not circularised. Nevertheless, the plasmid in the US genome is likely to be bigger than pRAS01 and pRAS02 based on the combined size of both contigs (Fig. S8b), and pRAS01 and pRAS02 may represent degraded variants of a larger plasmid. BLASTp analyses for the presence of RAGE genes revealed a complete Ti plasmid-like TraAI protein in the US plasmid contigs (Fig. S8b). Similar to pRAS02, F plasmid-like proteins were also not found in these contigs.

### Selective pressures on proteins of rickettsial vertebrate pathogens within the TRG

There is an increasing number of rickettsial species reported within the TRG, with some species known to infect vertebrates [87, 88]. We performed positive selection analysis using aBSREL with 492 single copy orthogroups (OGs) identified for the TRG (Fig. 8) to identify OGs under selective pressure among the vertebrate pathogens. A total of 105 OGs were identified with signals of positive selection in at least one branch at P < 0.05 after correction for multiple testing. Most of the OGs (88/105) with positive signals were observed only on the individual branches leading to the rickettsial species infecting *Ctenocephalides* spp. (*i*.*e*., *R. asembonensis* str. Perak, *Rickettsia* sp. TH2014, *R. felis* str. LSU) or the booklouse, *Liposcelis bostrychophila* (*R. felis* str. LSU Lb) (Fig. 8).

**Fig. 8.**
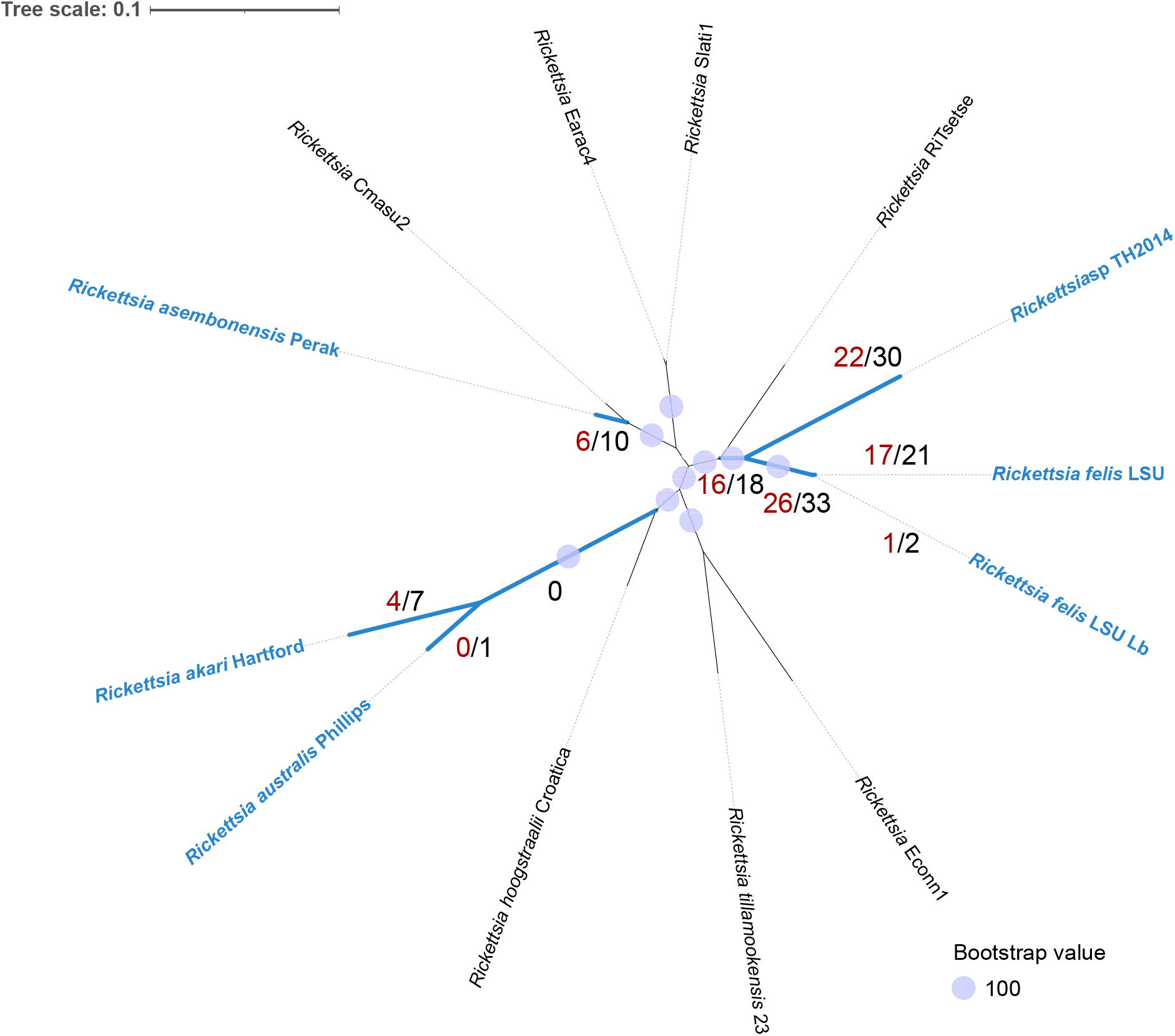
Maximum-likelihood tree based on 492 single copy orthogroups (OGs) identified for TRG rickettsiae (169,593 codon positions, 508,779 nucleotide positions) with best-fit model according to BIC: GY+F+R7. Codon-aware alignments of OGs were performed using PRANK. Branches marked blue were selected as foreground for aBSREL analysis of positive selection to identify OGs with signal of positive selection in at least one branch (P < 0.05 after correction for multiple testing). Red numbers adjacent to branches indicate the number of OGs with a signal of positive selection unique for that particular branch only. Black numbers indicate number of OGs with signals of positive selection at more than one branch. Species indicated in blue font can infect vertebrates.

The largest COG categories assigned to these OGs include translation (J – 15), energy production and conversion (C – 10), unknown function (S – 10), DNA replication and repair (L – 9) and cell wall/membrane/envelop biogenesis (M – 8) (Table S8). Several OGs comprised protein-coding genes predicted to be involved in lipopolysaccharide biosynthesis, peptidoglycan biosynthesis and outer membrane development, including *kdsB, lpxC, asmA, tolC*, and proteins with glycosyl transferase family domains. Moreover, protein-coding genes related to intracellular trafficking and secretion (*virB6, virB11, ftsY*) were also found to be under positive selection, alongside others involved in the ABC transport system (*lolD, rfbE, aprD* and *msbA2*), which could contribute to mechanisms of defence against antibiotics. Other protein coding genes with signals of positive selection that are potentially involved in vertebrate infection included *sca4* (on the *Rickettsia* sp. TH2014 branch) and a homolog of the ankyrin repeat protein RARP-1 (87.6% BLASTP percentage identity – on the *Rickettsia australis* branch, and the branch leading to both flea and booklouse *R. felis* strains).

### First complete genome of Bartonella clarridgeiae from Asia

Maximum-likelihood phylogeny revealed that the *Bartonella* sp. from the *C. orientis* metagenome clustered closely with *Bartonella clarridgeiae* 73 (accession: NC_014932.1) (Fig. 9a, Fig. S9a), bootstrap value 100). Comparison of the Malaysian strain to existing assemblies of reference strains (*B. clarridgeiae* 73 and ATCC 51734) showed high similarities and synteny (Fig. 9b,c). The genome of *B. clarridgeiae* from Malaysia (designated str. Perak) contained 1,249 and 1,336 CDS as annotated by Prokka and RASTtk (Fig. S9b), respectively, and 41 tRNA and six rRNA genes. One tmRNA gene was annotated by Prokka, and 40 repeat regions were annotated using RASTtk.

**Fig. 9.**
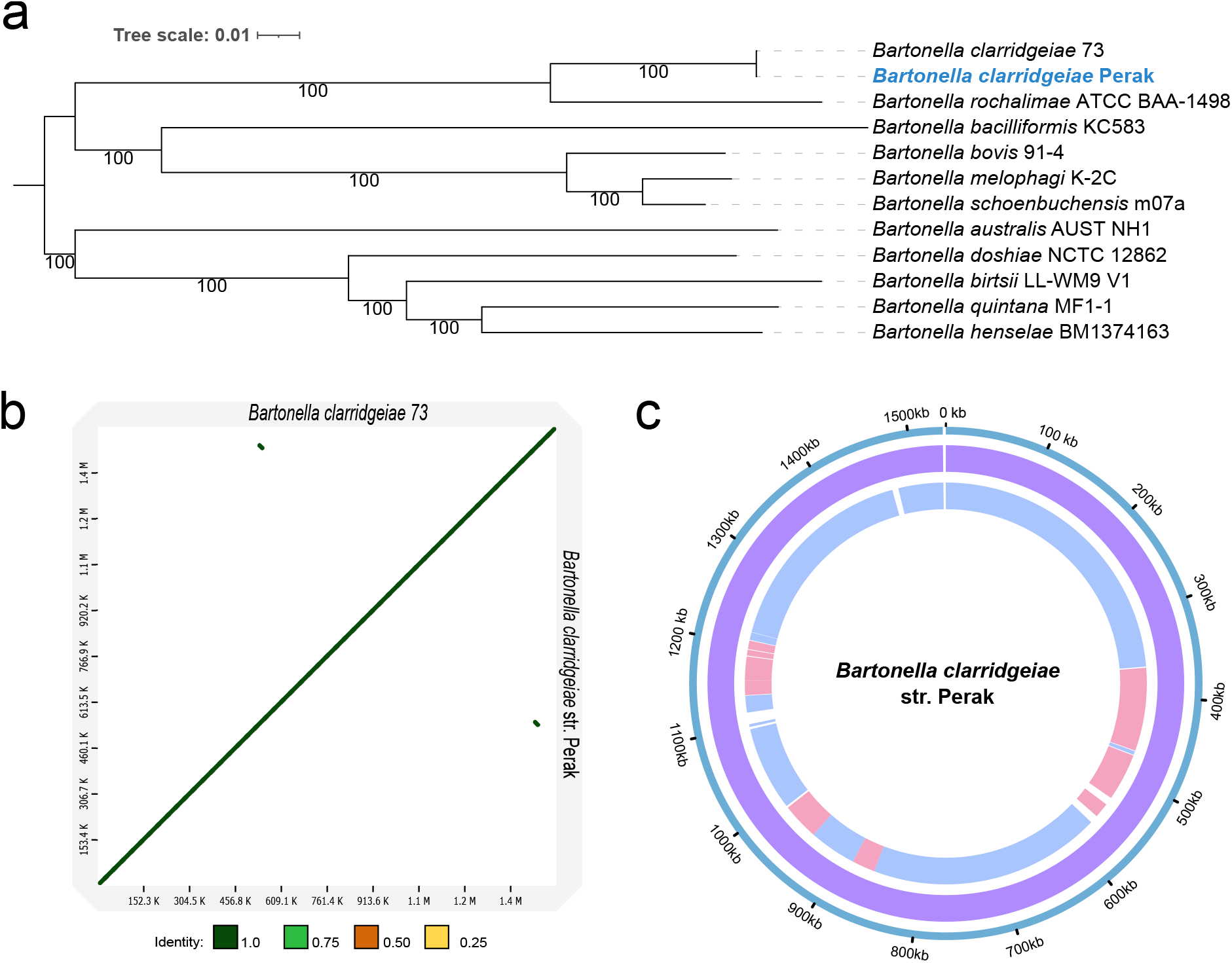
(a) Maximum-likelihood tree of selected *Bartonella* spp. based on 810 single copy orthogroups (OGs) (255,950 amino acid positions) using a partitioned best-fit model for each OG. Tree was rooted at mid-point. (b) Minimap2 alignment of *Bartonella clarridgeiae* str. Perak and *Bartonella clarridgeiae* 73 assemblies (accession: NC_014932.1). The identity values refer to a BLAST-like alignment identity computed from the alignment file and have been binned into four groups (<0.25, 0.25 – 0.5, 0.5 – 0.75, 0.75 – 1). (c) ProgressiveMauve alignments of *B. clarridgeiae* genomes. Outer ring: *B. clarridgeiae* str. Perak. First inner ring: locally colinear blocks identified between *B. clarridgeiae* str. Perak and *B. clarridgeiae* 73. Second inner ring: locally colinear blocks identified between *B. clarridgeiae* str. Perak and 19 contigs from the *B. clarridgeiae* ATCC 51734 assembly (accession: GCF_000518185.1).

### Variants in secretion, transport and effector proteins identified in B. clarridgeiae str. Perak

A total of 336 SNPs were identified when comparing the genome of *B. clarridgeaie* str. Perak with that of *B. clarridgeiae* 73 (Table S9), with 149 SNPs identified as non-synonymous variants. The majority of the non-synonymous variants (94) were identified within genes that encode proteins associated with secreted effector proteins and bacterial secretion systems (Table 2). A large proportion of the SNPs (76 non-synonymous, 108 synonymous) were located within a region of approximately 10 kb in length (positions 501,962 – 511,747 in the *B. clarridgeiae* 73 genome). This region consisted of several genes encoding proteins that constitute the flagellar secretion system and the inducible *Bartonella* autotransporter (iba) family protein. Three of the variants (one non-synonymous, two synonymous) were detected in positions corresponding to the variably expressed outer membrane protein (Vomp) family autotransporter/trimeric autotransporter adhesin gene. Sixteen synonymous substitutions, one frameshift variant and one deletion were detected in genes encoding for *Bartonella* effector proteins (Bep), the substrate of the VirB type IV secretion system (T4SS).

**Table 2.**
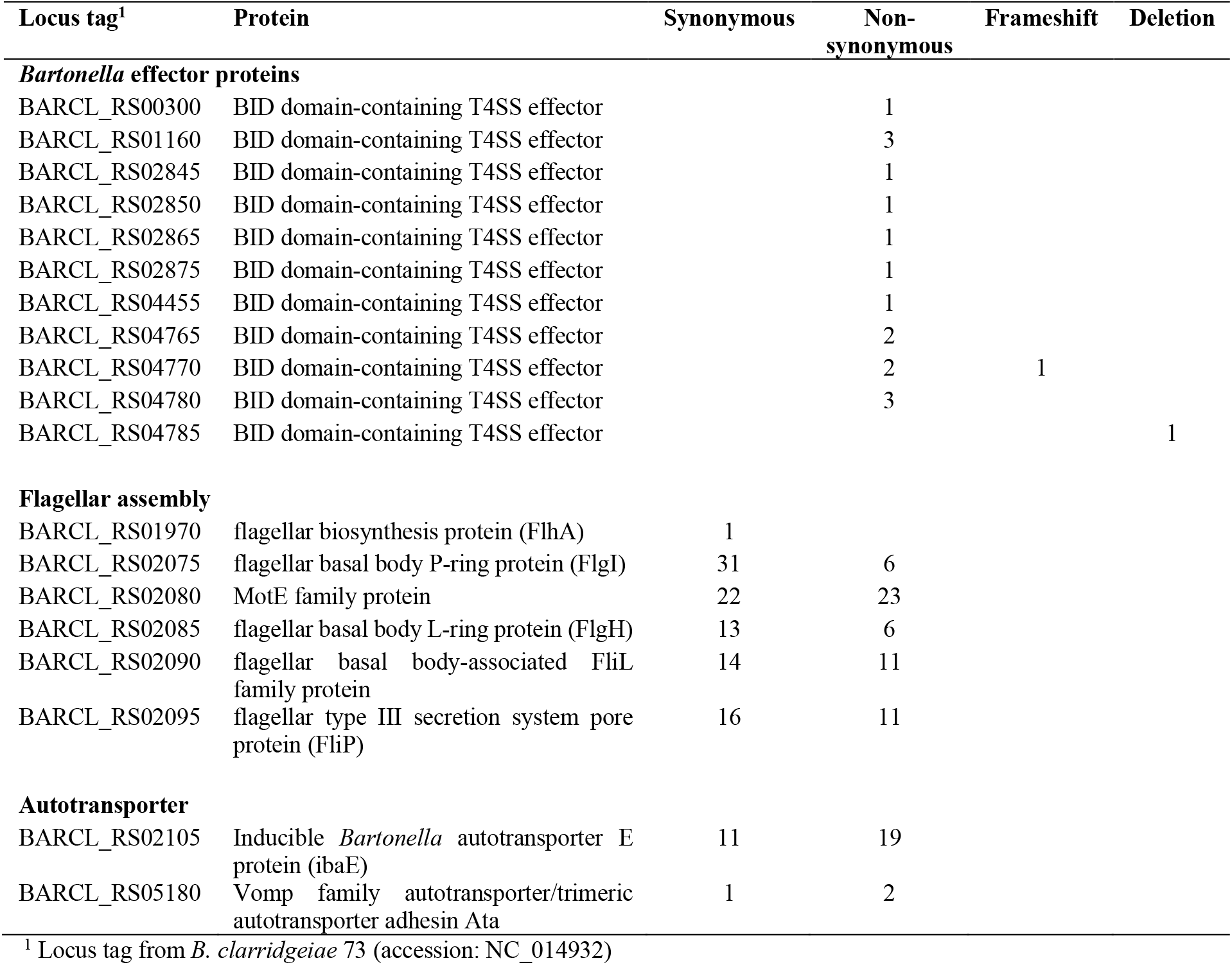
Selected genomic variants identified in *Bartonella clarridgeiae* str. Perak using *Bartonella clarridgeiae 73* as reference

### Mitochondrial genome assembly for C. orientis

Using the *C. felis* EL2017-DRISC mitogenome [89] as a template, we recovered three circularised assemblies for the *C. orientis* mitogenome; however, genome polishing was most effective for an assembly from sample pool F5 (Table 1). The assembly size was 22.19 Kb and annotation revealed all expected features except a lack of the origin of light strand replication (also absent in all other sequenced mitogenomes of Siphonaptera), while the origin of heavy strand replication was split into two pieces (Fig. S10a). We compared this assembly to the *C. felis* EL2017-DRISC mitogenome and a partial assembly of the *C. canis* mitogenome from China [90], which lacks most of the AT-rich control region (Fig. 11a). The three mitogenomes were highly similar to each other, with an extended control region apparent in *C. orientis* compared with *C. felis* EL2017-DRISC.

All mitochondrial assemblies of Siphonaptera shared the same complement of genes and synteny, although *Xenopsylla cheopis* harbours a fragmented paralog of the *nad2* gene (Fig. S10b). The *C. felis* record MT594468.1 is considered as a reference mitogenomic assembly, as it was obtained with PacBio technology, which in contrast with short-read technologies, usually resolves the long repeats of the control region without collapsing them. No information available regarding the sequencing technology used for the smaller *C. felis* assembly (MK941844) is available, but it was likely produced by a short-read method. For the single published *C. canis* mitogenomic assembly, the authors reported that the AT-loop region was incomplete, so the short control region is an artefact [90]. However, we achieved full resolution of the long control region in *C. orientis*, as some of the raw Nanopore reads spanned its entirety.

A phylogenomic analysis incorporating 15 mitochondrial genes revealed a well-supported, monophyletic clade for the family Pulicidae, with *C. orientis* exhibiting a closer relationship with *C. canis* than with *C. felis* (Fig. 10b). Although there are few complete mitogenomes for *C. felis* available, the two assemblies from China (MW420044 and MK941844) clustered separately from isolate EL2017-DRISC, which has an US origin (Fig. 10c). Concatenating two mitochondrial genes (partial *coi* and full-length *coii*) that are frequently used in flea phylogenetics enabled us to confirm the placement of our *C. orientis* sample within the clade formed by *C. orientis* voucher specimens (Fig. S11), all of which were of South-East Asian origin. The only unexpected finding was a *C. felis damarensis* specimen nested within the *C. orientis* branch. Recent morphological and molecular studies have indicated that this subspecies is of dubious validity [4, 91].

**Fig. 10.**
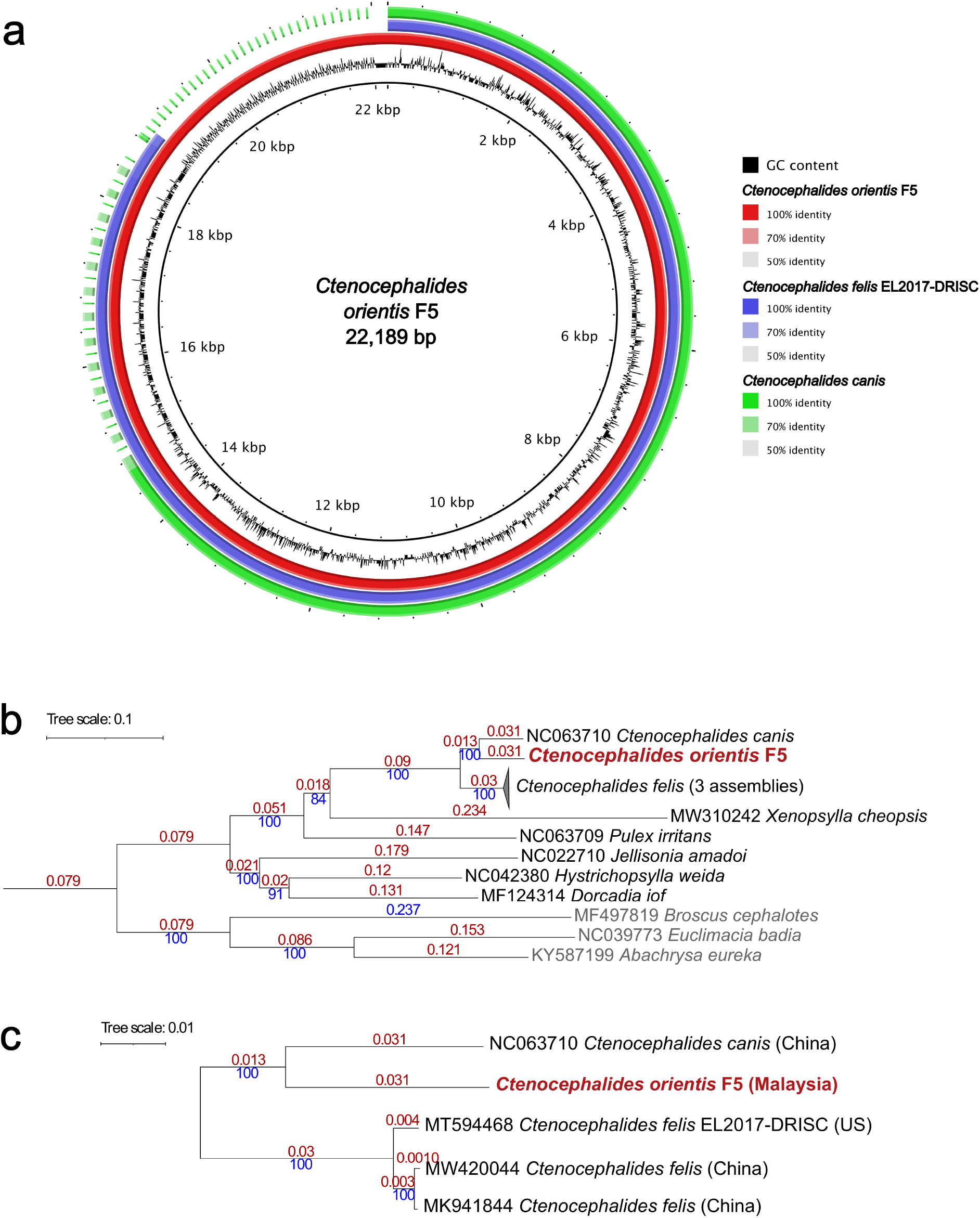
(a) A circular plot comparing three mitogenomic assemblies for *Ctenocephalides* spp.: *C. orientis* (F5, this study, accession: OQ383237), *C. felis* and *C. canis*. The intensity of the colour indicates the identity score of a BLAST alignment. (b) Maximum-likelihood tree based on concatenated alignments of 15 mitochondrial genes (Table S2, 12,476 nucleotide positions). Numbers marked with red represent branch length. Numbers marked with blue represent bootstrap support value. (c) Enlarged portion of the tree in (b) with the *C. felis* clade expanded.

## Discussion

Despite the cosmopolitan distribution of *Ctenocephalides* spp. fleas and their importance both as disease vectors and a cause of flea allergy dermatitis, a genome for a representative species (*C. felis*) only became available in 2020 [25] and few bacterial genomes associated with the *Ctenocephalides* spp. microbiome have been published to date. The present study provides new insights into the molecular taxonomy of *C. orientis* and the diversity and evolution of *Wolbachia* strains associated with *Ctenocephalides* spp. Moreover, it presents novel genomes for *R. asembonensis* and *B. clarridgeiae*, suspected and confirmed human pathogens respectively, for which genomic data were very sparse globally and entirely absent from Asia.

The “Oriental cat flea”, *C. orientis*, was originally considered as a subspecies of *C. felis*. However, over several decades, the identification of discrete morphological and molecular differences between *C. felis* and *C. orientis* have led to the elevation of the latter to full species status [92, 93]. A prior phylogenetic study based on two nuclear and two mitochondrial markers placed *C. orientis* and *C. canis* in a clade sister to *C. felis* [4], which was reproduced in the present study using 15 mitogenomic loci. Moreover, the natural history of *C. orientis* is clearly distinct from that of *C. felis*, as the former has a strong host preference for dogs [4], although it will also parasitize small ruminants [94]. Our discovery of a novel *Wolbachia* strain (*w*Cori) in *C. orientis* that displays important genomic differences compared to its counterpart in *C. felis* (*w*CfeT) suggests divergence in the microbiome of these two flea species, although systematic sampling across numerous geographic sites will be needed to determine if *w*Cori and *w*CfeT are specific to each flea host.

*Ctenocephalides felis* hosts the most diverse collection of *Wolbachia* strains for any arthropod species examined so far, with representatives of five supergroups (A, B, F, I and the proposed supergroup V) reported from different locations worldwide [28, 95] [26, 96-98]. The prevalence of the group F strain sequenced here in *C. felis* is unknown, but highly similar *Wolbachia* 16S rRNA gene sequences have been reported from *C. felis* collected in Georgia, USA [95] as well as in other regions of Malaysia [99]. When the group F culture isolate was first reported, the presence of a second *Wolbachia* strain in the tick cells used for intracellular culture was suspected due to the presence of double peaks in some of the Sanger-sequenced PCR products [28], but this was assumed to be closely related to the dominant group F symbiont. However, the metagenomic sequencing of the cultures performed here revealed the lower-density strain to be *w*CfeJ, first reported from a cat flea colony in California [26]. While *w*CfeJ is common in cat fleas from multiple laboratory colonies in the US, it is very rare in the wild according to sampling from the US and UK, with records from only two North American locations in the literature (Orange County, CA and Washington, NC) [17]. Thus, the presence of a highly similar *w*CfeJ isolate in Malaysia is remarkable. Here, we assign the American and Malaysian strains of *w*CfeJ to a new *Wolbachia* supergroup (V) that so far has no other members described worldwide, although it appears to have affinities with supergroup F, the *Wolbachia* clade with the broadest taxonomic host range [81].

Historically, the classification of *Wolbachia* phenotypes was shaped by comparison between the reproductive parasitism and evidence of horizontal transmission in supergroups A and B, and the apparent obligate mutualism and host-symbiont coevolution exhibited by the filarial nematode-specific clades C and D [100]. Although it is now known that *Wolbachia* co-evolution with filariae has broken down on multiple occasions, with numerous examples of symbiont loss and gain [101], evidence of co-evolution between *Wolbachia* and its hosts in some haematophagous arthropod taxa has provided new impetus to research on mutualistic relationships involving *Wolbachia* [82, 102, 103]. The best studied of these is the symbiosis between the bedbug *C. lectularius* and *Wolbachia* strain wCle from supergroup F, which is housed in a bacteriocyte and provisions riboflavin and biotin (deficient in blood diets), the latter via the BOOM [27]. The presence of BOOM in *w*CfeT led the authors of the *C. felis* genome study to hypothesize that this strain is also an obligate mutualist, but although it is widespread in *C. felis* populations in the US and UK, wild cat fleas lacking *Wolbachia* infections have been identified [25, 26]. While *w*Cori from *C. orientis* is clearly related to *w*CfeT, it lacks BOOM and also differs from *w*CfeT due to the presence of intact *cifA*/*B* homologues, suggesting a potential role in reproductive parasitism. The *C. felis* genome contains laterally transferred, CI-like antidote genes that could potentially neutralize *Wolbachia*-induced CI [26]; hence, determining if this is also true for *C. orientis* will be important to further our understanding of flea-*Wolbachia* relationships. The presence of pantothenate biosynthesis genes in these supergroup I genomes, which are otherwise very sparsely distributed in the accessory genome of *Wolbachia* [82], adds further complexity to the potential host-symbiont phenotypes mediated by *w*CfeT and *w*Cori. This may indicate that vitamin provisioning is important in the symbiosis of *Ctenocephalides* spp. with supergroup I, even if these *Wolbachia* strains are not universally present and they differ in capacity for biosynthesis of biotin.

The *w*CfeF strain, which also harbours BOOM, represents another example of a *Wolbachia* genome that combines signatures of potential reproductive parasitism and mutualism. However, on closer examination, the pseudogenisation in this strain of the biotin synthase gene (*bioB*), which is responsible for the conversion of dethiobiotin to biotin, is difficult to reconcile with a mutualistic role and suggests *w*CfeF represents an evolutionary snapshot of BOOM undergoing degradation. Indeed, partial or relic BOOM have been identified in the genomes of other *Wolbachia* strains and across the wider Rickettsiales [26, 27]. In most instances, BOOM components may serve primarily to reduce endobacterial dependence on host-derived biotin rather than underpinning a mutualistic relationship, with plasmids facilitating exchange both within the *Wolbachia* genus and between other obligate intracellular bacteria [26]. Since cat fleas and bedbugs are likely to be found together in human dwellings in some instances, horizontal transfer of *Wolbachia* strains and mobile elements between hosts could be facilitated by the free-living flea larvae, which feed on the faeces of adult fleas and environmental detritus (potentially contaminated with the faeces of bedbugs and other arthropods). In support of this hypothesis, exchange of *Wolbachia* strains through a common environmental medium has been documented in several other contexts [104-107].

The *w*CfeF genome exhibited two other remarkable features that are unique to supergroup F: intact prophage regions and the presence of *cifA/cifB* homologues. Thus, alongside supergroups A, B and V, *w*Cori (supergroup I) and *w*CfeF significantly extend the range of *Wolbachia* clades that could potentially induce CI, although experimental validation that type V *cifA*/*cifB* homologues are functional in this respect is needed. Taken together, the genomics of *Ctenocephalides* spp.-associated *Wolbachia* reflect an unparalleled hotbed of *Wolbachia* evolution involving extensive LGT but also a geographically stable, albeit rare, strain in the form of *w*CfeJ [17].

*Rickettsia asembonensis* was first isolated in culture in 2015 [13] and formally described as a new species the following year [108], but sequences now considered as *R. asembonensis* had been identified in fleas from the Thai-Myanmar border in 2003 [109] and subsequently from a number of other locations worldwide [15]. The organism was known originally only from fleas and other arthropods until 2016, when sequences similar to those obtained from the Thai-Myanmar border study were amplified from the blood of a febrile child in Malaysia [110]. Following this report, *R. asembonensis* DNA has been detected in samples from febrile patients in Peru [111] and in two human cases in Zambia [112], although no categoric link with symptomatic disease was possible in the latter study. Moreover, *R. asembonensis*-like DNA sequences have been obtained from the blood of non-human mammals, including macaques [113] and domestic ruminants [114] in Malaysia. The *R. asembonensis* genome reported in the present study is the first complete, circularized genome from this emerging pathogen worldwide and was obtained from a country where zoonotic spill-over to humans is an ongoing risk. Notably, an isolate of *Ca*. R. senegalensis (*Rickettsia* sp. TH2014) [14] was obtained from *C. orientis* in the same region of Perak where *C. orientis* was collected for the present study, indicating that both rickettsial species are co-circulating in this vector in Malaysia, as is the case for *C. felis* in many other countries [17, 115-117]. The pathogenicity of *Ca*. R. senegalensis for humans is unclear, but evidence of infection in cats has been obtained from the US [118].

Although the *R. asembonensis* str. Perak chromosome was very similar to that of the only other sequenced strain (NMRCii from Kenya [108]), substantial differences in the plasmid structure and gene content between the strains were apparent. Our analysis of cat flea metagenomic data from the US [39] revealed the presence of a plasmid that had not been recognized, since the authors of the US study mapped their data to the NMRCii reference genome and pRAS01 is not closely related to pRAS02. These findings suggest that the plasmidome of *R. asembonensis* is undergoing evolutionary change on different continents while the chromosome remains remarkably stable. However, the *R. asembonensis* plasmids are cryptic, containing neither the active RAGE and BOOM present in the plasmids of the tick symbiont *R. buchneri* [84], nor the intact conjugative apparatus of the plasmids of *R. felis* [6]. A recent genus-wide analysis of the rickettsial plasmidome showed that over a quarter of the gene content is not found in the pan-chromosomal gene repertoire and has no homologs in other organisms [88]. Rickettsial plasmids were postulated to be vertically inherited from a common ancestor and were present in 56% of the 34 species included in Karkouri *et al*.’s analysis [88] (33% of 80 strains they studied). While plasmids are frequently lost in *Rickettsia* spp., there were very few cases identified where plasmid gene content was present in the chromosome of plasmid-free strains [88], suggesting that rickettsial plasmids may consist mostly of selfish elements.

Classically, rickettsial pathogens were classified in the typhus group (vectored by fleas or lice) and the spotted fever group (vectored mainly by ticks). Subsequently, *R. bellii* and *R. canadensis* were placed in their own “ancestral” clades, while the genomic characterization of *R. felis* led to the establishment of the TRG [6], which includes close relatives of *R. felis* transmitted predominantly by fleas, as well as other pathogens transmitted by ticks (*R. australis*) or mites (*R. akari*). However, with the explosion of metagenomic datasets available for invertebrates other than arthropod vectors, it is now recognised that most rickettsial diversity is harboured by non-haematophagous arthropods and various aquatic invertebrates [87]. The majority of these rickettsial symbionts are classified in the *Torix* group (named after a genus of leeches [119]) but some of the exclusively invertebrate-associated *Rickettsia* spp. belong to the TRG [87]. Thus, as the term “transitional” suggests, the TRG represents a bridge between the diversity of rickettsial symbionts vertically transmitted within a single invertebrate host species and those that have adapted to a lifecycle including a vertebrate host. Interestingly, recent data have suggested that a booklouse-associated strain of *R. felis* may cause disease in humans via airborne dispersal, despite being found in a non-ectoparasitic host [120].

Our analysis of positive selection on the branches of the TRG leading to vertebrate pathogens identified a number of genes that may have a key role in adaptation to the mammalian cellular niche and evasion of the more sophisticated immune response of vertebrates, including genes involved in LPS synthesis and outer membrane structure. Two genes under selection on some of the branches associated with infection of vertebrates, *sca4* and *rarp-1*, were of particular interest, as their role in the cell biology of *Rickettsia parkeri* (a member of the SFG) has recently been experimentally investigated. Sca4 is a secreted effector that mediates spread between host cells by facilitating protrusion formation through interactions with host vinculin, which reduces intercellular tension [121]. In contrast, RARP-1 is a periplasmic ankyrin protein that is not secreted into host cells but nevertheless contributes to rickettsial entry, perhaps via regulation of type IV secretion system components [122]. However, not every branch of the phylogeny involving infection of vertebrates exhibited evidence of selection within these two genes.

The final component of the *C. orientis* microbiome sequenced here, *B. clarridgeiae*, has been detected frequently in *Ctenocephalides* spp. fleas and the blood or other tissues of cats and dogs worldwide [17, 123-125], with a number of reports across South-East Asia [126-129], although to the best of our knowledge no confirmed zoonotic infections from this part of the world have been documented. However, only two other genomes are available (one complete and circularised [130]), which are derived from feline blood isolates from France and the USA. Thus, the *B. clarridgeiae* genome presented here is the first to be sequenced directly from a flea, the first associated with a canine host, and the first from Asia. Relatively few SNPs were detected compared with the feline isolates and it was striking that the vast majority of these were concentrated in regions involved in host cell entry, especially the flagellum, secretion systems and Beps.

*Bartonella clarridgeiae* is a member of lineage 3, in which the flagellum is critical for invasion of host erythrocytes [131]. In contrast, the VirB/D4 type IV secretion system has a key role in establishment of the intracellular niche during infection of nucleated cells (likely to be predominantly endothelial cells) in the primary phase of infection preceding erythrocyte colonisation [132]. The Beps that are secreted via the type IV secretion system inhibit endocytic uptake of *Bartonella*, instead facilitating the formation of an “invasome” [133], in which large aggregations of bacteria are internalised into the host cell. The Beps are also important in inhibition of apoptosis and subversion of intracellular signalling, although their specific roles in lineage 3 have been little studied [134].

Additional SNPs in the Perak strain were also apparent in the iba and trimeric autotransporter adhesin (TAA) gene, which belong to the type Vc secretion system [135]. The best-studied TAA in *Bartonella* spp. is BadA of *B. henselae*, which forms pilus-like structures on the bacterial surface responsible for autoagglutination between bartonellae, attachment to the extracellular matrix of the host, targeting of β1-integrins on endothelial cells, and pro-angiogenesis [136]. In other bartonellae, Vomps may play a similar role to BadA. These immunodominant proteins undergo recombination in *Bartonella quintana* and inactivation of one or more members has been observed in different strains of this species, although loss of all four Vomps resulted in avirulence [137]. A similar process of inactivation may occur in *B. clarridgeiae*, as missense SNPs was detected in the TAA gene. Less is known about the function of iba genes in bartonellae, but they are upregulated during infection of endothelial cells and one member of this group may have cohemolysin (erythrocyte membrane disruption) activity in *B. henselae* [138]. Taken together, the concentration of SNPs in protein-coding genes directly involved in establishment of infection may reflect local adaptation to the *C. orientis*-dog life-cycle compared with the reference genome from a cat, but unfortunately *B. clarridgeiae* is too little studied, both experimentally and genomically, for this hypothesis to be further evaluated at this time.

In conclusion, our metagenomic analysis of *Ctenocephalides* spp. homogenates and bacterial isolates highlights a picture of relative genome stability in the key three components of the microbiome (*Wolbachia, Rickettsia* and *Bartonella*), although signatures of evolution were apparent in genes involved in interactions with the flea or the vertebrate host. Unlike most other obligate haematophagous arthropods [139], there is no evidence for a primary endosymbiont in fleas that provisions vitamins, although some *Wolbachia* strains may partake in this role as secondary symbionts. Our study also provides important insights into two poorly-characterised, cosmopolitan zoonotic pathogens, which showed limited geographic variation except in mobile elements (*R. asembonensis*) and genes required for entry into mammalian cells (*B. clarridgeiae*). Moreover, over longer timescales, genes involved in cell invasion were also found to be under selection in the flea-associated rickettsiae of the TRG. Finally, this study showed that our strategy to isolate rickettsial agents from *Ctenocephalides* spp. fleas using tick cell lines and the metagenomic sequencing of whole flea samples facilitated the detection and characterisation of other symbionts such as *Wolbachia*. Further studies are needed to understand the basis of differences in virulence between *R. felis, R. asembonensis* and *Ca*. R. senegalensis, to obtain pure cultures of the two *Wolbachia* strains *w*CfeF and *w*CfeJ, and to elucidate whether *Wolbachia* infections interact positively or negatively with zoonotic pathogens vectored by fleas [140].

## Author statements

### Author contributions

Conceptualization, S.A., A.C.D., B.L.M. and J.J.K.; methodology, A.B., K.K.T., A.S., L.B.S., A.C.D., B.L.M., and J.J.K.; formal analysis, A.B., K.K.T., A.S., A.C.D., B.L.M., and J.J.K.; investigation, A.B., K.K.T., A.S., N.A.H., F.S.L., L.B.S., and J.J.K.; resources, L.B.S., C.K.S.C., S.A., A.C.D., and B.L.M.; data curation, A.B., N.A.H. and J.J.K.; writing—original draft preparation, A.B., K.K.T., B.L.M., and J.J.K.; writing—review and editing, A.B., K.K.T., A.S., S.K.L., A.S., C.K.S.C., S.A., A.C.D., B.L.M. and J.J.K.; visualization, A.B., K.K.T., A.S. and J.J.K.; supervision, L.B.S., C.K.S.C., S.A., A.C.D., and B.L.M.; project administration, S.K.L. and J.J.K..; funding acquisition, L.B.S., S.A., A.C.D., B.L.M., and J.J.K.

### Ethical approval

This research was conducted as part of a larger surveillance study with the approval from the Department of Orang Asli Development [JHEOA.PP.30.052 Jld. 6 (19)] and the Universiti Malaya Institutional Animal Care and Use Committee (G8/23122019/11102019-01/R).

### Consent for publication

Not applicable

## Acknowledgements

The authors would like to thank the Orang Asli community at Kampung Tumbuh Hangat, Perak for their assistance in the fieldwork. The tick cell lines were provided by the Tick Cell Biobank Asia Outpost at Universiti Malaya and the Tick Cell Biobank at the University of Liverpool. The IDE8 cell line was used by kind permission of Prof. Ulrike Munderloh, University of Minnesota.

