## Supplementary figures for "Metagenomics of culture isolates and insect tissue illuminate the evolution of *Wolbachia, Rickettsia* and *Bartonella* symbionts in *Ctenocephalides* spp. fleas"

a

| Contig | Length | Coverage | Circular |
|---|---|---|---|
| **contig_1** | 1451344 | 55 | Y |
| **contig_2** | 1199327 | 25 | Y |

b

| Contig | Length | Coverage | Circular |
|---|---|---|---|
| **contig_1** | 1450987 | 38 | N |
| **contig_2** | 1191286 | 16 | N |

c

| Contig | Length | Coverage | Circular |
|---|---|---|---|
| **contig_1** | 1201492 | 64 | Y |
| **contig_2** | 1451057 | 108 | Y |

d

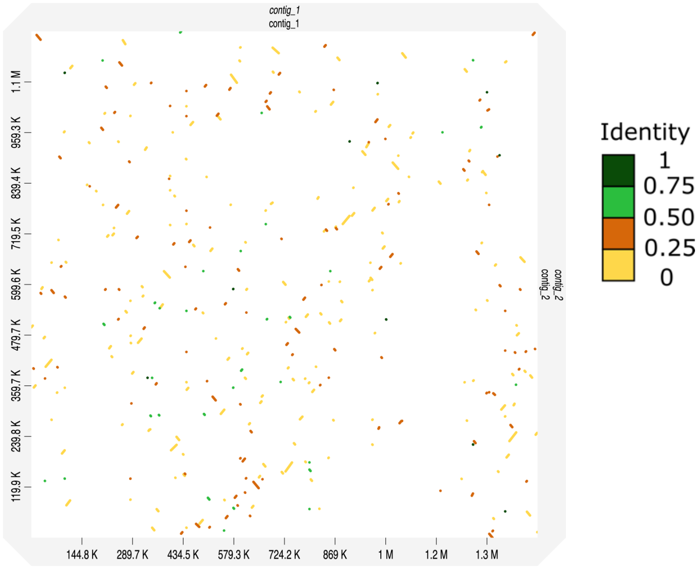


Fig. S1. (a, b and c) Flye assembly output for Nanopore sequencing reads from (a) *Wolbachia*-infected IDE8 parent culture, (b) *Wolbachia*-infected IDE8 P4 subculture and (c) *Wolbachia*-infected BME/CTVM23 P1 subculture.
(d) Dotplot generated from Minimap2 alignment of contig_1 (horizontal, ~1.4 mb) and contig_2 (vertical, ~1.2 mb) generated from the Flye assembly of Nanopore sequencing reads from *Wolbachia*-infected IDE8 parent culture. The identity values refer to a BLAST-like alignment identity computed from the alignment file and have been binned into four groups (<0.25, 0.25 – 0.5, 0.5 – 0.75, 0.75 – 1).

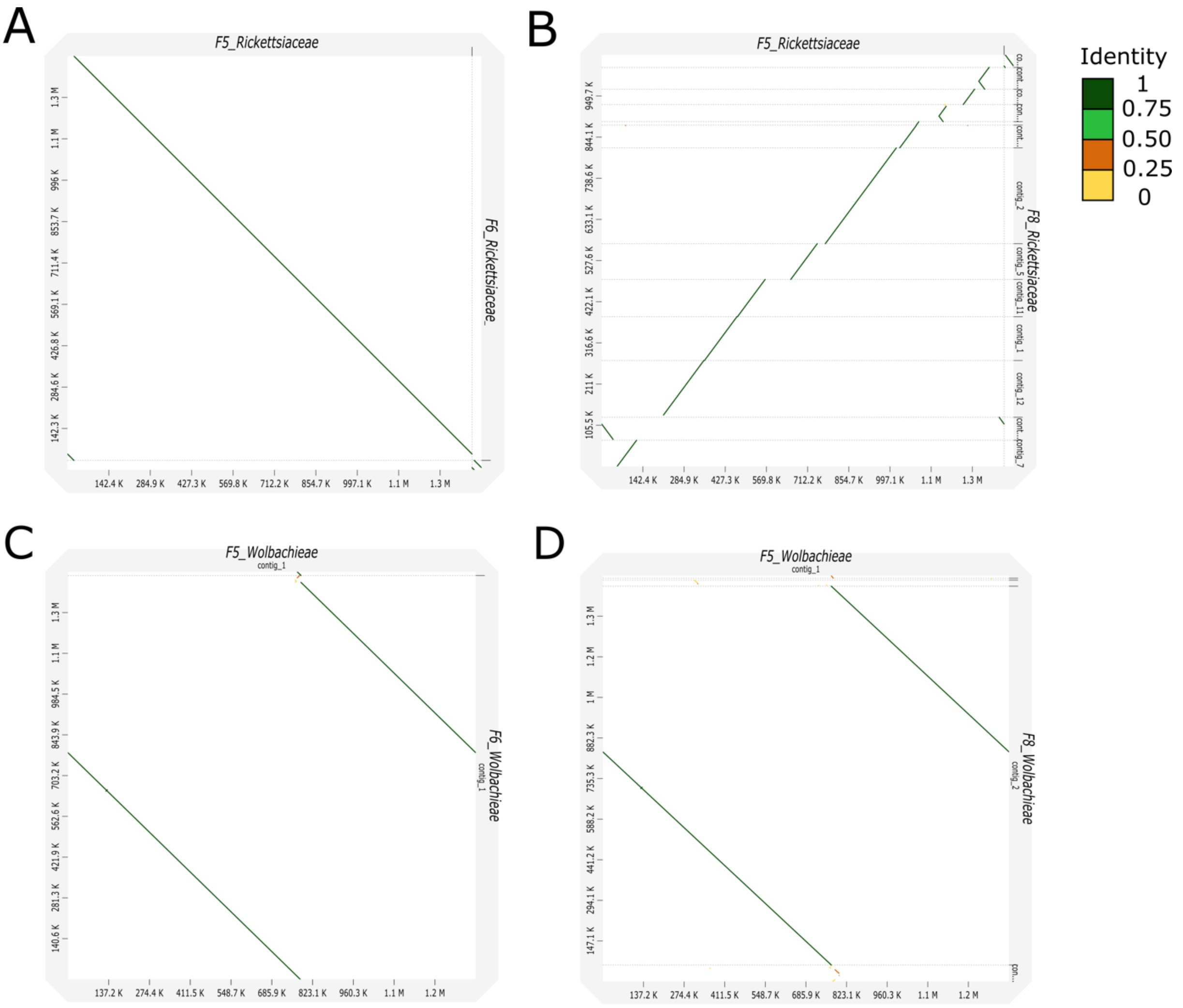


Fig. S2. Dotplot generated from Minimap2 alignment of *Rickettsiaceae* and *Wolbachia* (*Wolbachieae*) Flye assemblies from *Ctenocephalides orientis* pools. A and B: *Rickettsiaceae* assemblies from F5 (horizontal axis) and F6 (A, vertical axis) or F8 (B, vertical axis). C and D: *Wolbachia* Flye assemblies from F5 (horizontal axis) and F6 (A, vertical axis) or F8 (B, vertical axis). The identity values refer to a BLAST-like alignment identity computed from the alignment file and have been binned into four groups (<0.25, 0.25 – 0.5, 0.5 – 0.75, 0.75 – 1).

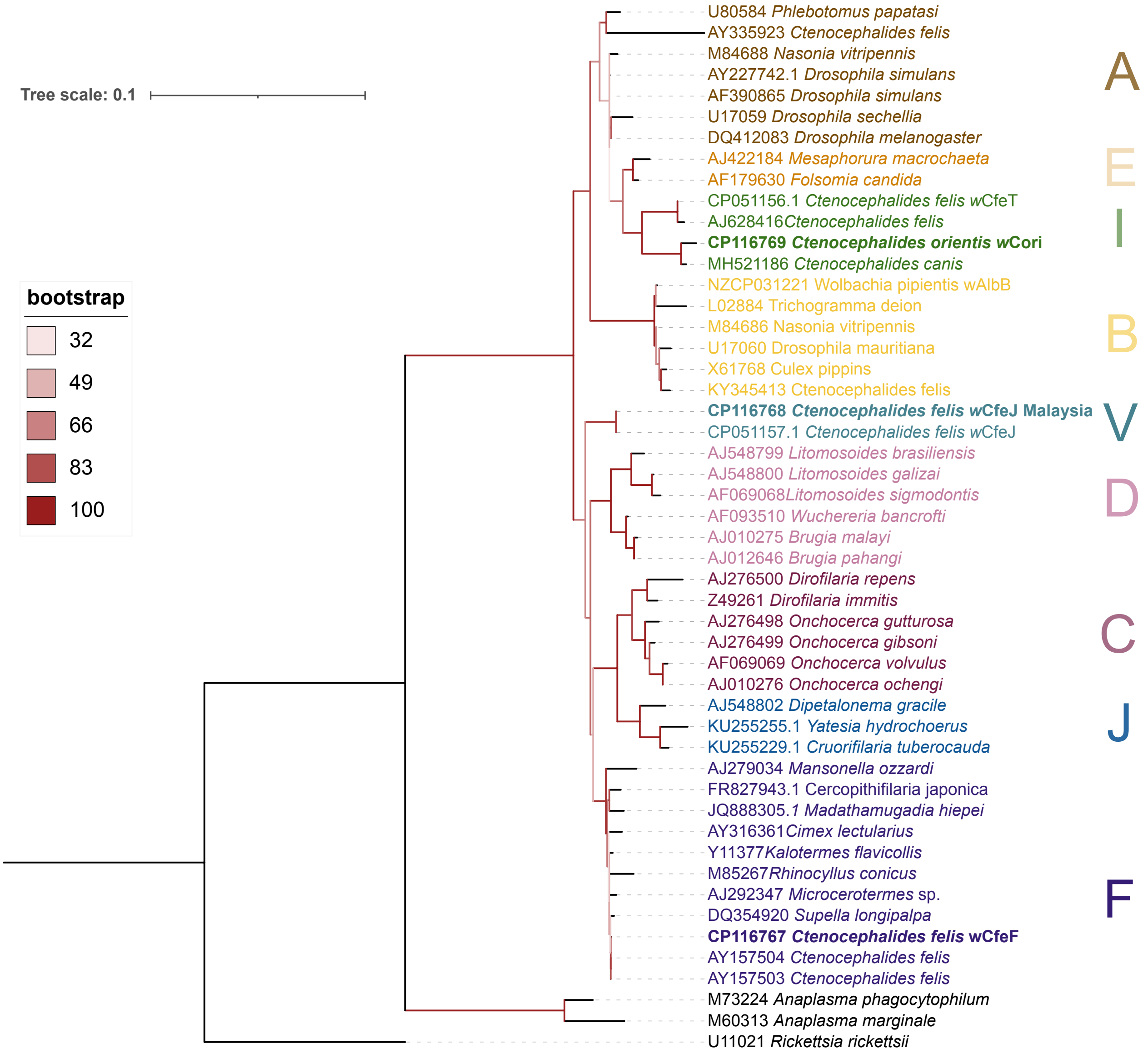


Fig. S3. Maximum likelihood phylogeny based on partial 16S rRNA sequences (1,615 nucleotide positions, best model according to BIC: HKY+F+R2) of *Wolbachia* strains from the indicated hosts. Accession numbers are given for the sequences in the NCBI GenBank database. Other members of the Rickettsiales, *Rickettsia rickettsii*, *Anaplasma phagocytophilum* and *Anaplasma marginale*, were used as an outgroup

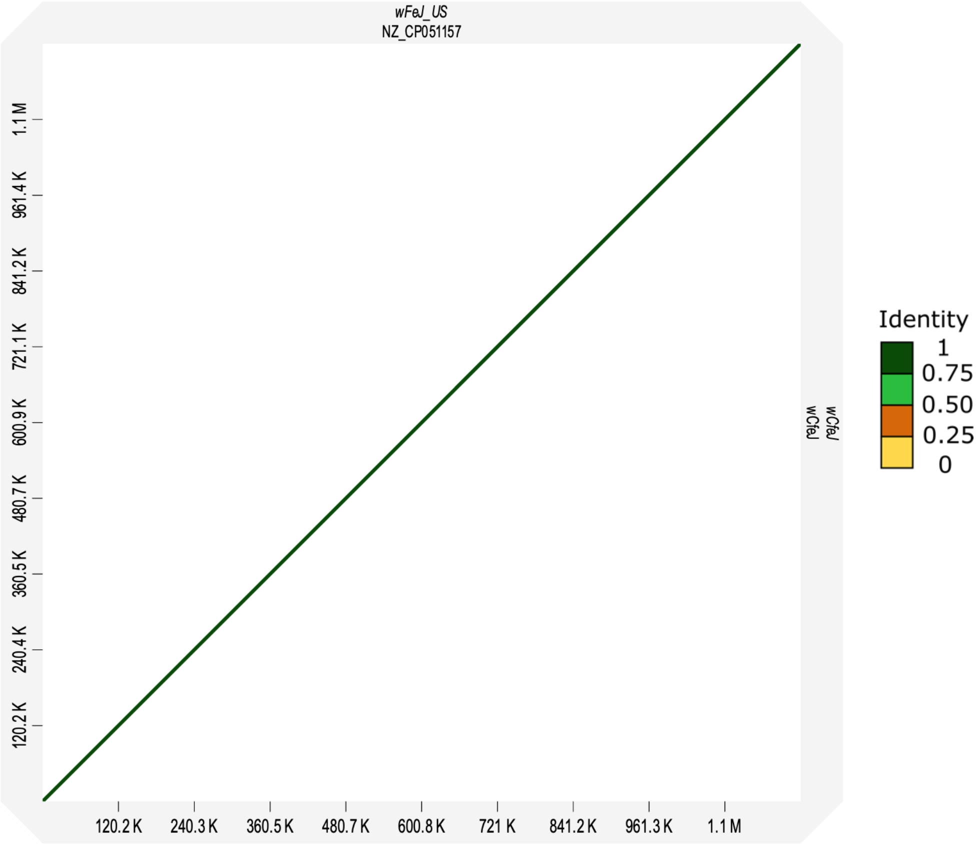


Fig. S4. Minimap2 alignment of *w*CfeJ from US (horizontal, Accession: NZ_CP051157.1) and *w*CfeJ isolate from the current study (vertical,) The identity values refer to a BLAST-like alignment identity computed from the alignment file and have been binned into four groups (<0.25, 0.25 – 0.5, 0.5 – 0.75, 0.75 – 1).

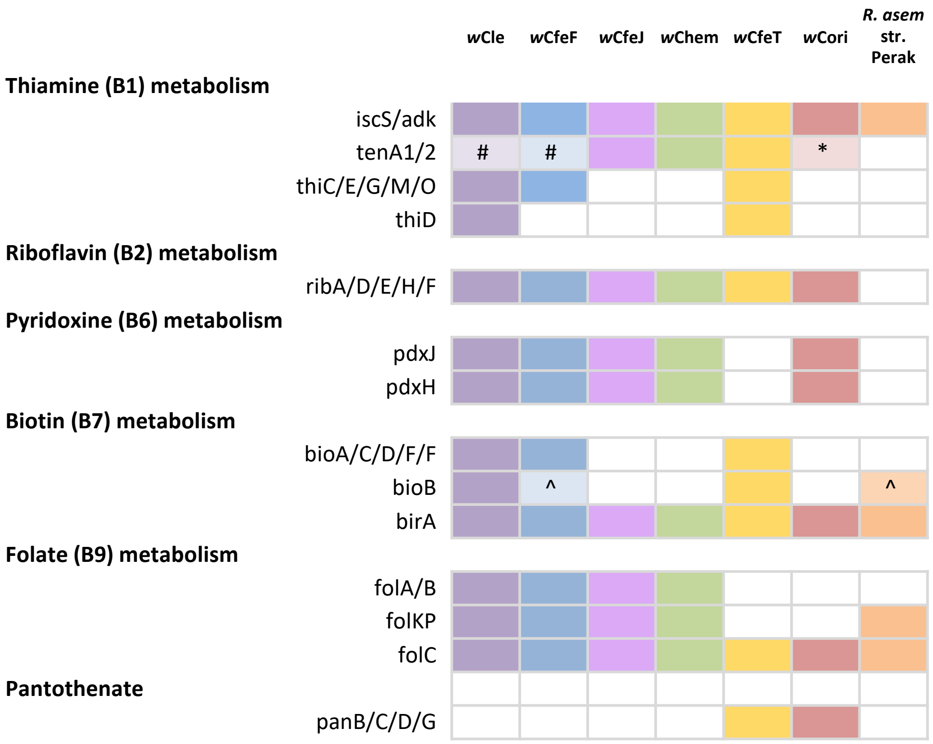


### only one copy
* truncated
^ fragmented
@incomplete

Fig. S5. KEGG functional analyses of genome data obtained from *Wolbachia* strains *w*Cle, *w*CFeF, *w*CfeJ, *w*Chem, *w*CfeT and *w*Cori and *Rickettsia asembonensis* str. Perak from the present study and published datasets.

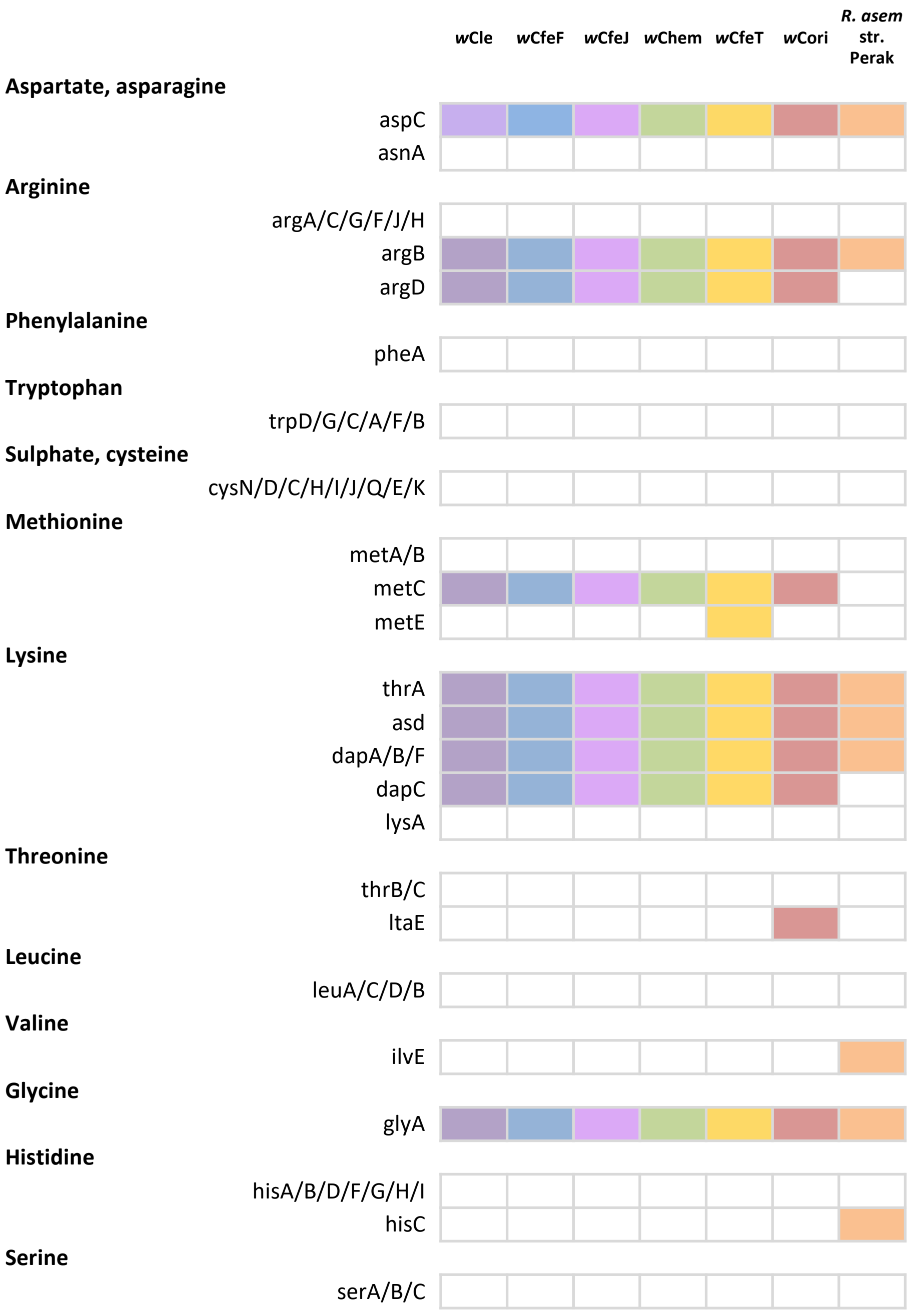


Fig. S5. KEGG functional analyses of *Wolbachia* and *Rickettsia*. (continued from previous page)

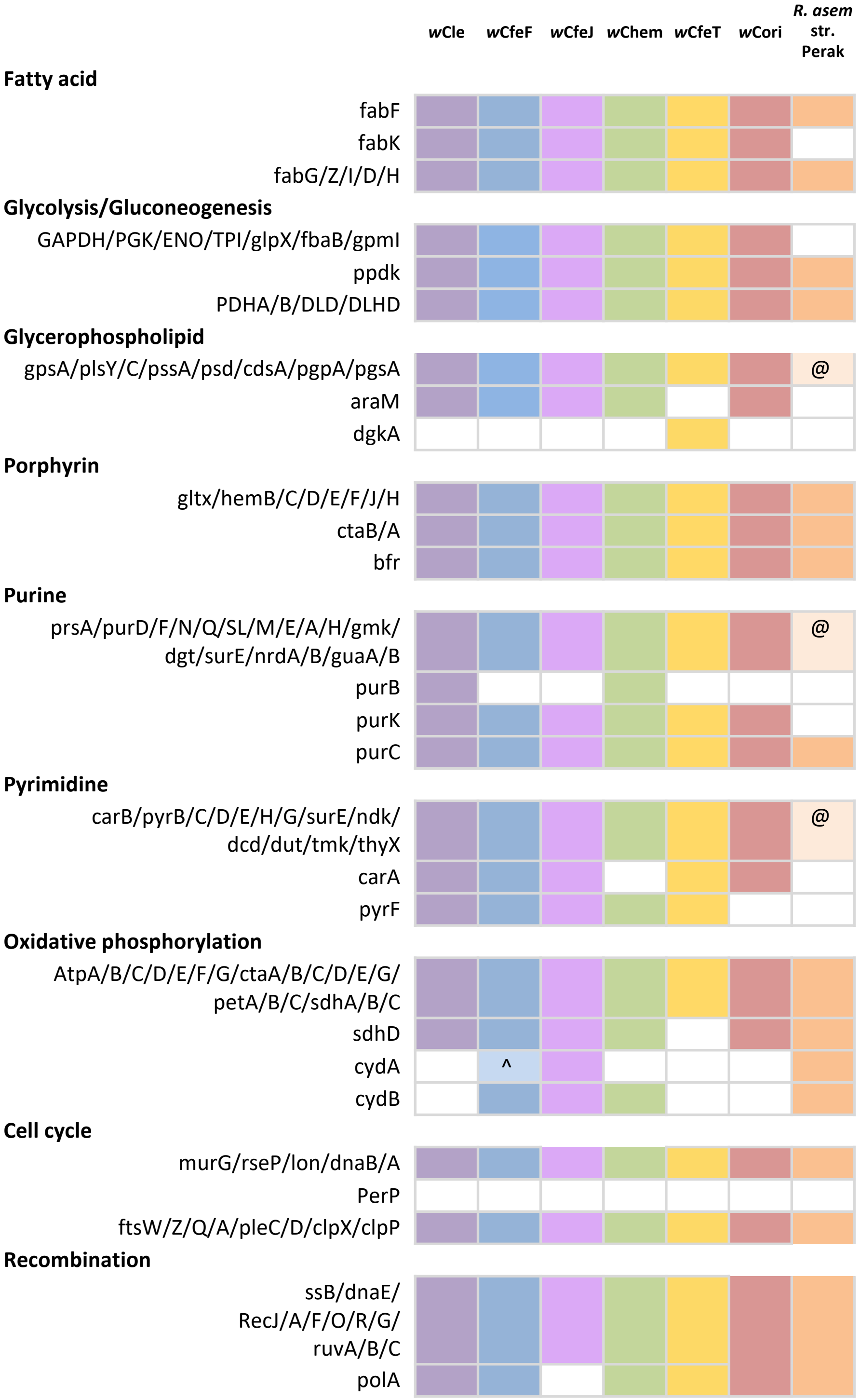


Fig. S5. KEGG functional analyses of *Wolbachia* and *Rickettsia*. (continued from previous page)

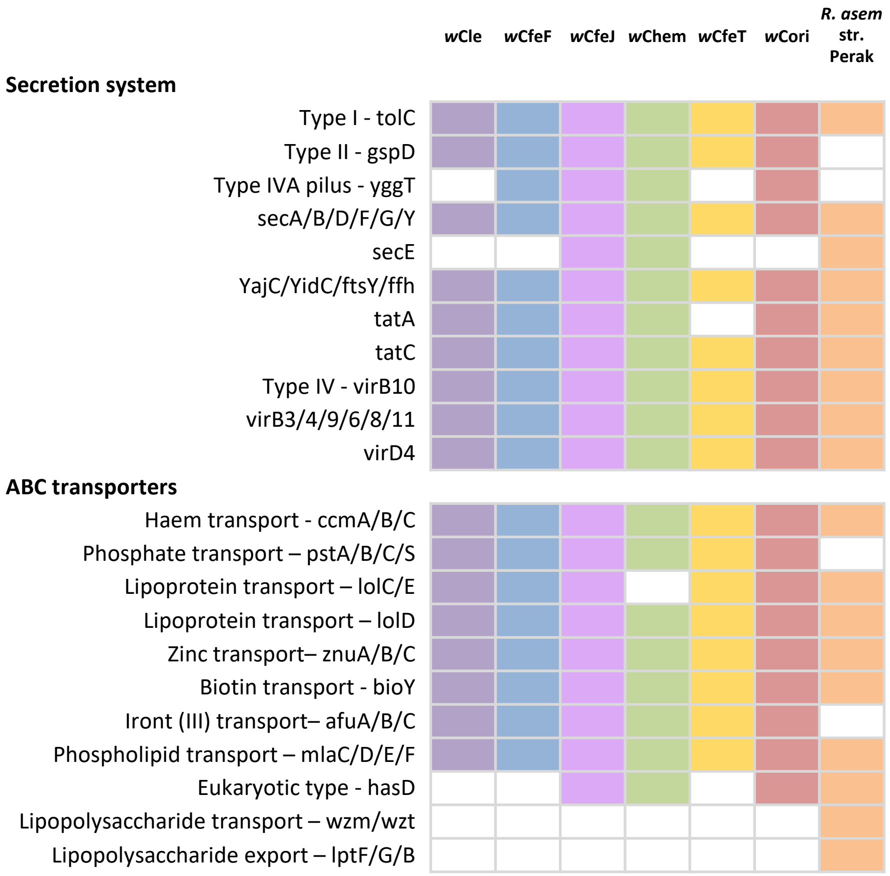


Fig. S5. KEGG functional analyses of *Wolbachia* and *Rickettsia*. (continued from previous page)

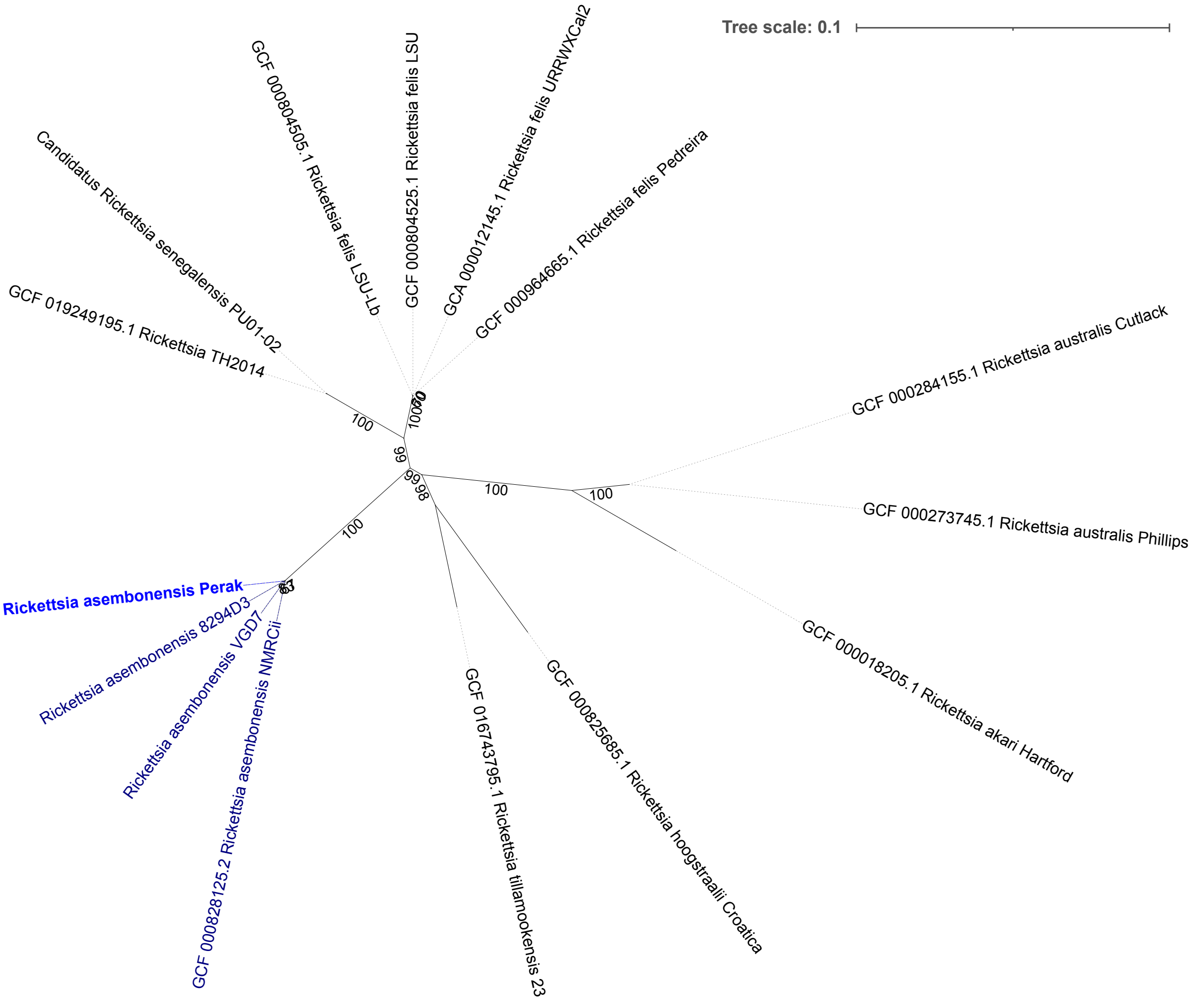


Fig. S6. Maximum-likelihood tree of selected *Rickettsia* spp. based on three separate genes, *gltA*, *sca4* and *ompB* (9,587 nucleotide positions) with best-fit model according to BIC: TVM+F+G4.

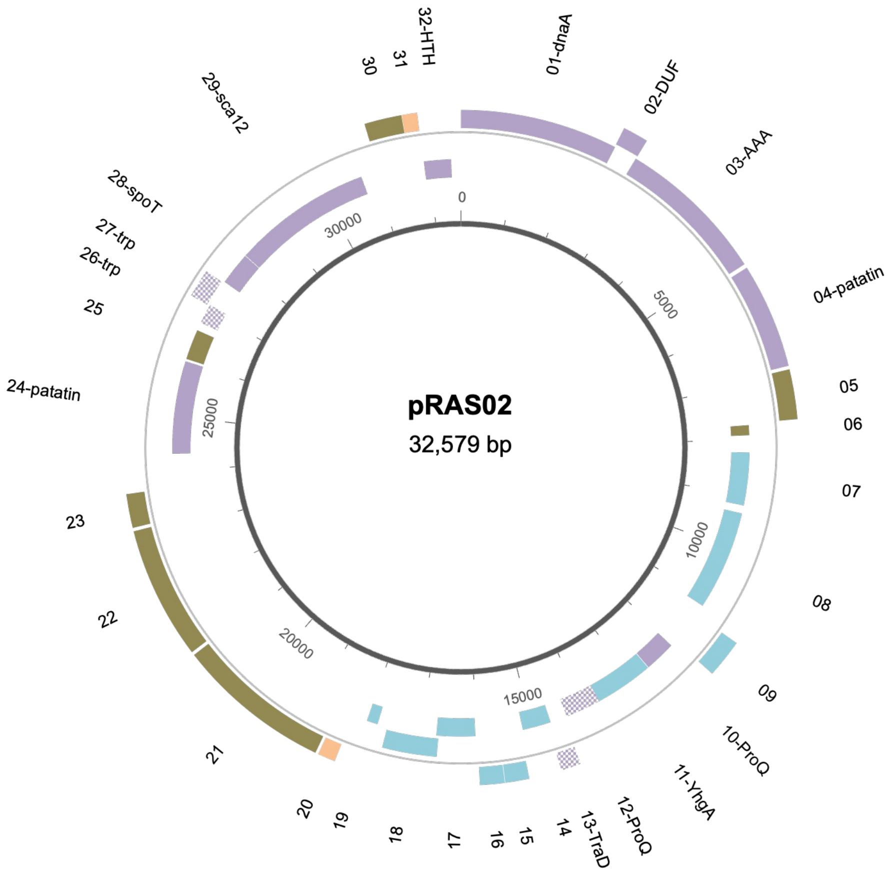


| | Matched proteins in BlastP | Percentage ID | Evalue |
|---|---|---|---|
| 01 | dnaA-like | 54.83% | 0 |
| 02 | DUF1828 | 73.83% | 3E-49 |
| 03 | AAA family ATPase | 70.89% | 0 |
| 04 | patatin-like phospholipase | 93.77% | 0 |
| 10 | ProQ activator of osmoprotectant ProP | 76.44% | 7.00E-89 |
| 12 | ProQ/FinQ family protein | 76.89% | 1.00E-106 |
| 13 | Putative conjugative transfer protein traD | 100.00% | 2.00E-59 |
| 24 | patatin-like phospholipase | 80.24% | 0 |
| 26 | tetratricopeptide repeat protein | 64.67% | 1.00E-52 |
| 27 | tetratricopeptide repeat protein | 37.32% | 2.00E-14 |
| 28 | spoT bifunctional (p)ppGpp synthetase | 85.28% | 1.00E-121 |
| 29 | autotransporter domain-containing protein | 80.77% | 0 |
| 32 | helix-turn-helix transcriptional regulator | 39.75% | 4.00E-33 |

Fig. S7. Map of *Rickettsia asembonensis* plasmid pRAS02. CDS are shown on forward strand outside grey line and the reverse inside the grey line. CDS were colour-coded as the following: CDS encoding proteins with Blastp matches to other known proteins (purple), CDS encoding transposase, insertion sequences, and recombinases (teal), CDS encoding hypothetical proteins, CDS encoding proteins with no Blastp matches (orange). Chequered boxes represent fragmented CDS.

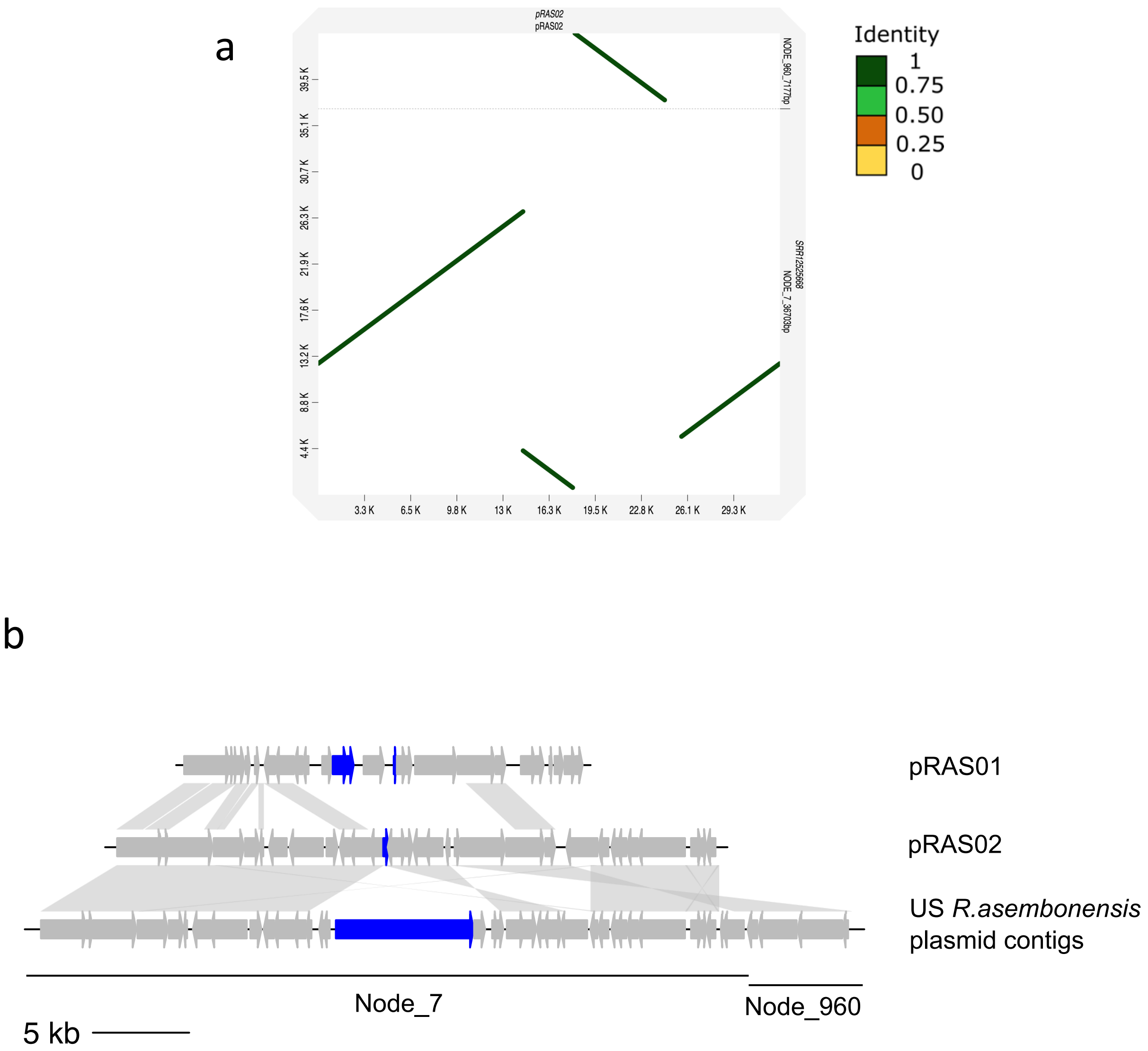


Fig. S8. (a) Minimap2 alignment of *Rickettsia asembonensis* plasmid pRAS02 (horizontal axis) with two contigs identified from the US *R. asembonensis* assembly (Node_960 and Node_7). The identity values refer to a BLAST-like alignment identity computed from the alignment file and have been binned into four groups (<0.25, 0.25 – 0.5, 0.5 – 0.75, 0.75 – 1). (b) BLASTn comparison of pRAS01 (*R. asembonensis* str. NMRCii), pRAS02 (*R. asembonensis* str. Perak) and the concatenated contigs identified from the US *R. asembonensis* assembly. Assemblies were rotated to start at *dnaA*-like gene. Blue colour denotes the Ti plasmid-like genes.

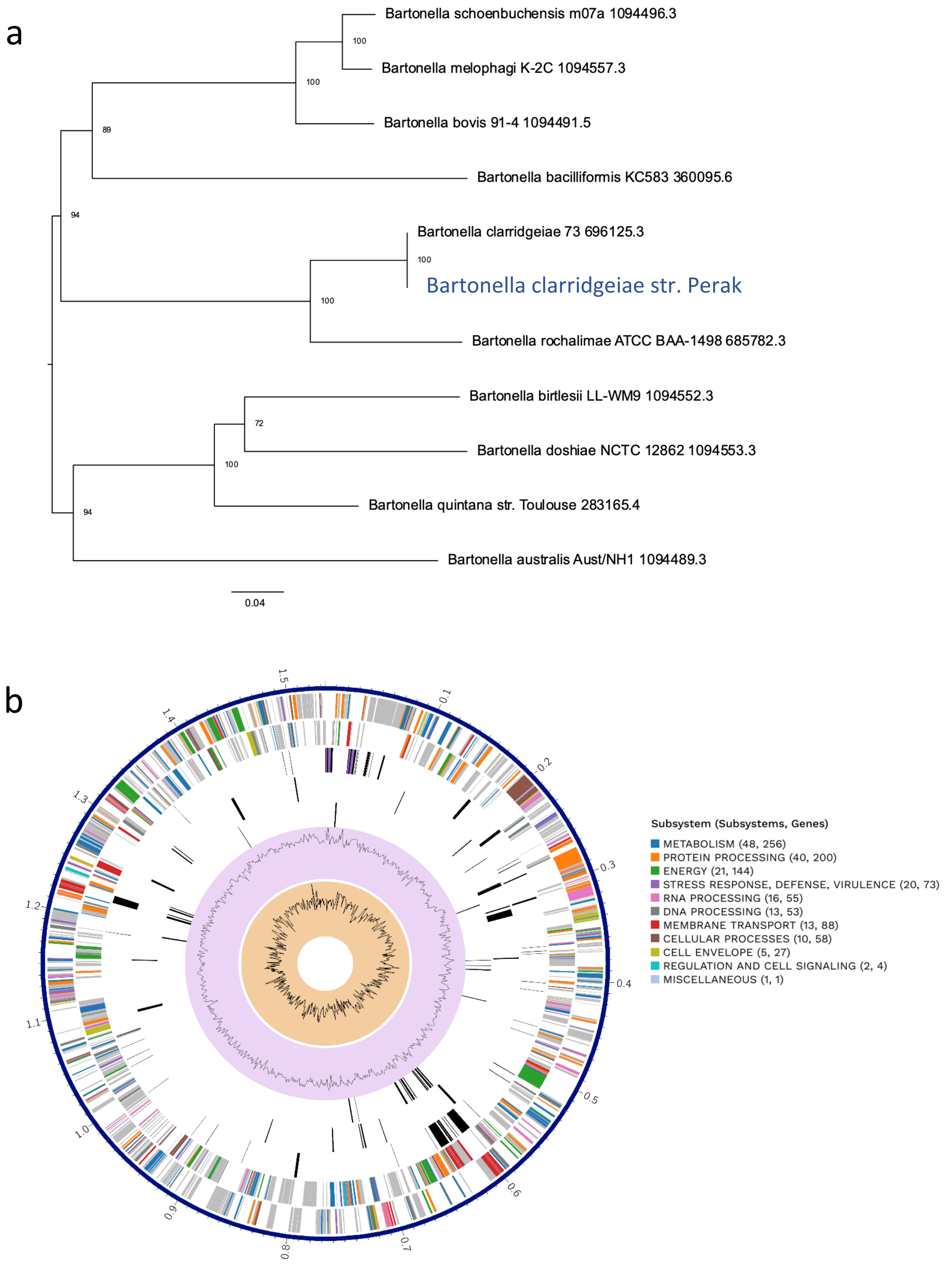


Fig. S9. Phylogeny of *Bartonella* spp. and annotation of *Bartonella clarridgeiae* str. Perak from the the PATRIC comprehensive genome analysis pipeline. (a) RAxML tree of *Bartonella* spp. based on 708 single copy genes (762,406 positions). (b) Circos representation of *Bartonella clarridgeiae* str. Perak genome. The genome annotations were displayed from outer to inner rings: 1) CDS on the forward strand, 2) CDS on the reverse strand, 3) RNA genes, 4) CDS with homology to known antimicrobial resistance genes, 5) CDS with homology to known virulence factors, 6) GC content, and 7) GC skew.

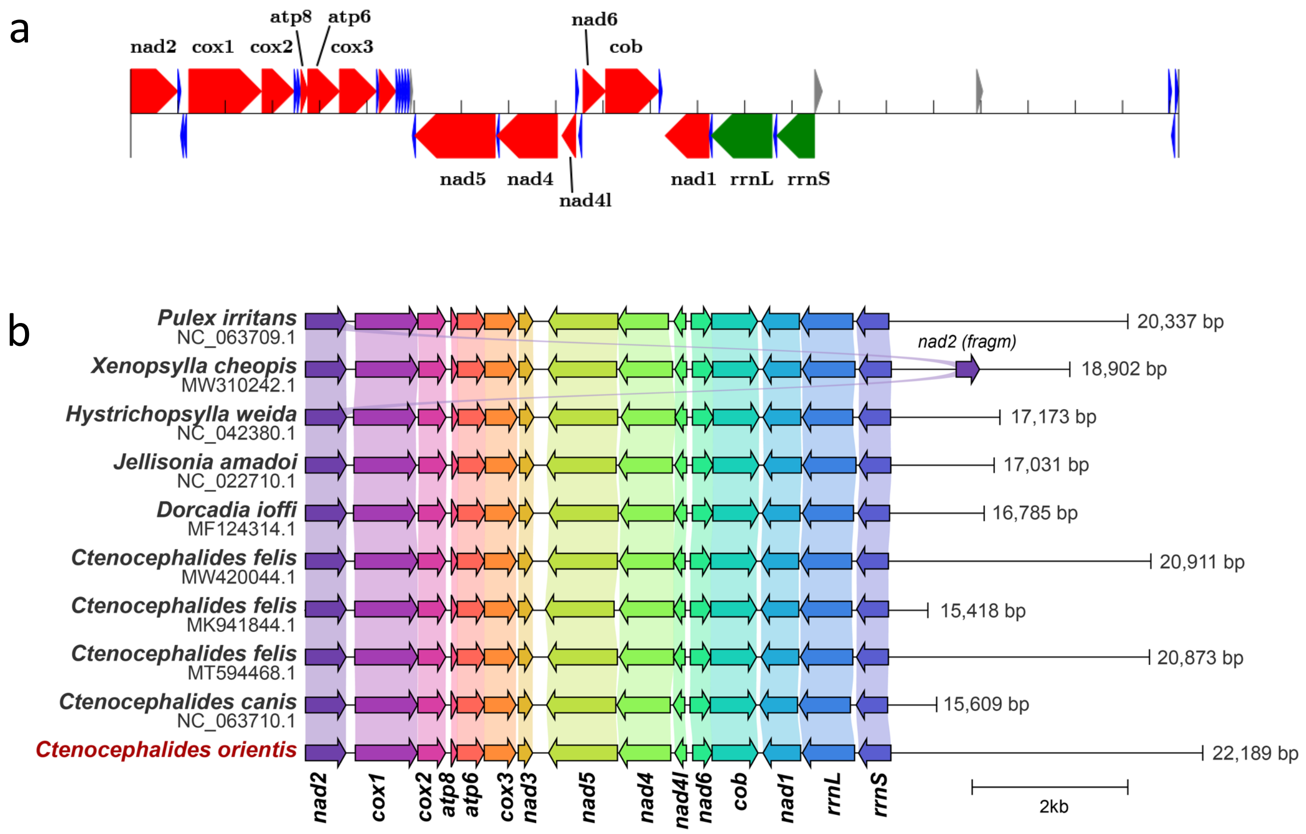


Fig. S10. (a) A map of a mitochondrial assembly of *Ctenocephalides orientis*. Green arrows represent ribosomal genes, red – protein coding genes, blue – tRNA genes, grey – OH (origin of heavy strand replication). (b) Linearised mitochondrial gene arrangement within *Siphonaptera* species. Gene and genome sizes are to scale. No tRNA genes nor origin of replication are present in this scheme.

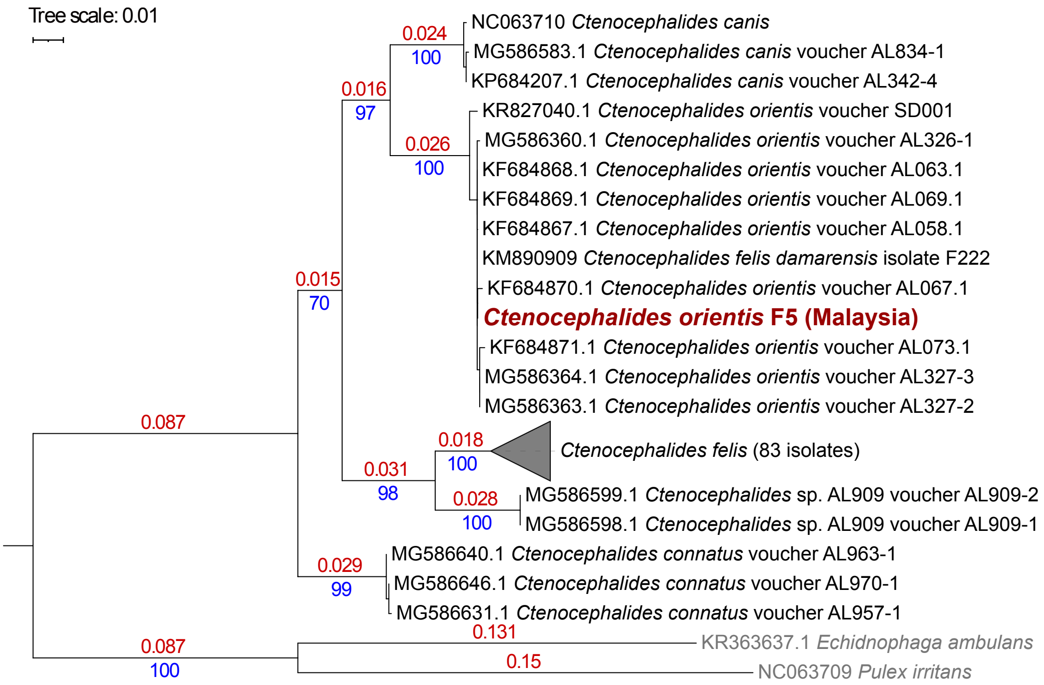


Fig. S11. Maximum-likelihood tree based on concatenated alignments of partial *coi* and full-length *coii* sequences (1,328 nucleotide positions total) with best-fit model according to BIC: TPM1uf+G4 for both genes. Numbers marked with red represent branch length. Numbers marked with blue represent bootstrap support value. Sequences for *Ctenocephalides orientis* (Malaysia) used in this study were obtained from mitochondrial genome assembled from F5 pool (Table 1 – accession: OQ383237 ).
